# Cryptic selection forces and dynamic heritability in generalized phenotypic evolution

**DOI:** 10.1101/291963

**Authors:** William Gilpin, Marcus W. Feldman

**Affiliations:** Department of Applied Physics; Department of Biology, Stanford University, Stanford, CA

## Abstract

Individuals with different phenotypes can have widely-varying responses to natural selection, yet many classical approaches to evolutionary dynamics emphasize how a population’s average phenotype increases in fitness over time. However, recent experimental results have produced examples of populations that have multiple fitness peaks, or that experience frequency-dependence that affects the direction and strength of selection on certain individuals. Here, we extend classical fitness gradient formulations of natural selection in order to describe the dynamics of a phenotype distribution in terms of its moments—such as the mean, variance, skewness, etc. The number of governing equations in our model can be adjusted in order to capture different degrees of detail about the population. We compare our simplified model to direct Wright-Fisher simulations of evolution in several canonical fitness landscapes, we find that our model provides a low-dimensional description of complex dynamics not typically explained by classical theory, such as cryptic selection forces due to selection on trait ranges, time-variation of the heritability, and nonlinear responses to stabilizing or disruptive selection due to asymmetric trait distributions. In addition to providing a framework for extending general understanding of common qualitative concepts in phenotypic evolution—such as fitness gradients, selection pressures, and heritability—our approach has practical importance for studying evolution in contexts in which genetic analysis is infeasible.

## 1 Introduction

The effects of evolutionary forces may be apparent in natural populations even when their underlying genetic consequences are not known. The size of river guppies increases when their natural predators are depleted;^1^ the beaks of Darwin’s finches grow larger after droughts;^2^ and mammals grow smaller in response to climate change.^3^ These and other natural and experimental studies demonstrate that rapid selection can produce noticeable changes in specific traits, underscoring the importance of considering phenotypic models of natural selection. These models are particularly relevant to studies in the field or of the fossil record where genetic analysis is unavailable or infeasible.^4^

A widely-used framework for such theories is the fitness landscape,^5^ an abstract function that describes the collective survival or reproductive benefits conferred by a given phenotype: an evolving population typically approaches a locally or globally maximum value in this space, subject to constraints on its rate of adaptation. But the underlying dynamics of this process may depend strongly on the context,^6^ and molecular techniques have only recently begun to shed light on the individual steps of adaptation and the intermediate phenotypes they produce.^7, 8^ Moreover, macroscale analyses have produced examples of non-monotone fitness functions with elaborate, multipeaked topographies subject to strong frequency dependence,^9^ selection that acts on trait ranges in addition to average values,^10^ and selection forces with nonlinear effects on metric traits.^11^ These studies indicate the need for simple, analytical models that can provide heuristic insight into the complex dynamics of phenotypic adaptation. Of particular importance are cases in which traditional experimental metrics, such as the realized heritability or selection response, have complex time-dependence over long timescales,^12, 13^ rendering derived experimental quantities such as selection coefficients insufficient as large-scale descriptors of population dynamics.

Many classical approaches to phenotypic evolutionary theory readily describes the evolution of a bell-shaped and fixed-width trait distribution within a given fitness landscape, resulting in the widely-known result that the rate of adaptation of the mean trait is directly proportional to the gradient of the mean fitness.^14–17^ This depiction of evolution as a fitness gradient-climbing problem is particularly intuitive, and it mirrors earlier work on the adaptive landscape of individual genes.^5^ However, many phenotypes fail to satisfy these conditions,^10, 18, 19^ as a result of which many experimental and theoretical studies have noted strong limitations on the applicability of fitness gradient dynamics.^20–23^ Efforts to establish more general rules for the evolution of an arbitrary trait distribution typically reformulate the underlying mathematics in terms of agent-based rules or a stochastic transmission kernel;^24–26^ however, these alternative formulations are difficult to compare directly with classical fitness gradient dynamics, and they typically introduce new assumptions regarding the underlying genetic processes or functional forms of the trait distribution or fitness.

Here, we develop a general model of phenotypic evolution that seeks to avoid imposing functional constraints on the fitness or trait by using a moment series expansion of the fitness landscape. Our approach has its origin in classical approaches that describe evolution of the mean trait as a fitness gradient-climbing problem, but we add additional dynamical equations for the variance, skewness, kurtosis, and finer-scale statistical features of the trait distribution. By adding or removing dynamical equations that describe various moments of the trait distribution, the level of detail our model captures about the evolutionary dynamics may be tuned. Importantly, our model reduces to a classical fitness gradient model when only the mean trait is allowed to vary. Our model explicitly relates the topology of the fitness landscape to the timescales (and thus dynamical relevance) of various phenomena through a series of coupling constants, which we compute analytically for several canonical fitness landscapes in a series of demonstrative examples. Using these examples, we show how even a simple generalization of classical fitness gradient dynamics can lead to a series of surprising effects in the evolutionary dynamics, including cryptic forces of selection that cause changes in the trait distribution even when the local fitness gradient is zero, as well as suppression of disruptive selection due to asymmetry and skewness in the trait distribution. We also show that our model can be re-formulated in terms of the coupled dynamics of the narrow-sense heritability and the mean fitness, and we show how these two quantities may jointly evolve under various conditions.

## 2 Model

Our model is based on a series expansion of the trait distribution in terms of its moments (such as the mean trait, trait variance, trait skewness, etc). Each moment has a separate dynamical equation, which couples its dynamics to those of the remaining moments. The strength and magnitude of the coupling of each moment’s dynamics to other moments depends on the specific fitness landscape, and can be summarized in terms of a finite set of “coupling coefficients” intrinsic to the landscape. Our basic approach is shown schematically in Figure S1, and below we describe how we derive dynamical equations for the mean trait 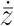, trait range 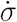, and mean fitness 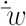. We also describe below the assumptions of our approach, as well as the relationship between our approach and similar series-based approaches developed by others.

## 2.1 Trait mean dynamics in an arbitrary phenotype distribution

We follow the classical approach to deriving phenotypic evolution originally developed by Lande.^15^ We start with the breeder’s equation, which relates the dynamics of the trait mean *z* to its change after a period of selection

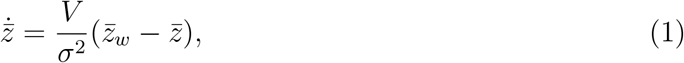

where the mean trait value 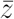, mean fitness 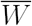, and mean trait after selection 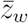 are defined in terms of statistical averages over the entire population,

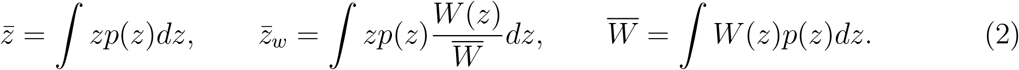

Here *p*(*z*) is the frequency of the trait *z* in the population, *W*(*z*) is an individual’s fitness as a function of its trait. The prefactor *V/σ*^2^ in Eq. 1 is equivalent to the narrow-sense heritability *h*^2^ of the trait, with *V* representing the additive genetic variance. Eq. 1 incorporates the primary assumptions regarding the underlying genetics; namely, that there is no direct gene-environment interaction, and that there is a linear regression between the selection differential and the mean trait value over short timescales^27, 28^ (although the slope of this regression, *h*^2^, need not be constant). In many previous studies in which Eq. 1 appears, the phenotypic variance *σ*^2^ is also assumed to be fixed, either due to logarithmic ranges in the values of metric traits that suppress the magnitude of fluctuations in trait variance, or due to the assumption of a fixed trait distribution shape that becomes a Gaussian distribution after an appropriate nonlinear transformation.^15, 27^ We will relax this requirement here.

Constancy of genetic variance *V* must be decided empirically for a given system under study; in general, *V* may change due to drift or direct gene-environment interactions.^13, 28, 29^ However, in the model of phenotypic evolution presented here, as in similar approaches based on the dynamics of single continuous traits,^30–32^ *V* may be assumed to vary much more slowly than the phenotypic variance *σ*^2^ provided there is linkage equilibrium, weak selection, or a near-equilibrium trait distribution. Additionally, for cases in which *V* changes quickly, such as during periods of strong directional selection or from transient effects such as the accumulation of linkage disequilibrium, rapid compensatory effects (for example, recombination) can allow *V* to stabilize at a well-defined average value over long timescales.^1, 12^ Accordingly, low or stationary genetic variance has been reported in certain experimental systems that may satisfy these conditions.^33^

Here, we use a standardized trait distribution 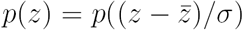 and also parametrize the fitness as 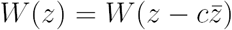, where *c* is an arbitrary positive constant. The case *c* = 0 corresponds to a trait that directly affects survival independently of other individuals in the population (examples include metabolic efficiency, camouflage coloration, or immune system competency), while the case *c =* 1 corresponds to a trait that affects fitness of a given individual only relative to others in the population (examples include secondary sex characteristics or running speed relative to a herd). Under this assumption, taking the derivative of both sides of Eq. 2 with respect to 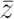 yields a simple relationship between an individual’s fitness landscape and the population mean fitness,

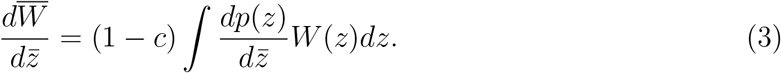

See Appendix C for full derivation. We note that if *c =* 1, then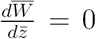; thus when the fitness landscape depends only on the difference between each individual trait and the trait mean, the mean fitness itself cannot depend on the mean trait.

Next, we assume that the trait distribution before selection *p*(*z*) is separable into a Gaussian part 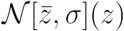 and a dimensionless non-Gaussian component *f*, which without loss of generality we express in terms of the dimensionless standardized coordinate 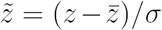,

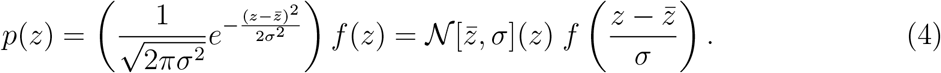

We emphasize that *f* = 1 corresponds to a purely Gaussian trait distribution, in which case our theory recreates standard evolutionary dynamics.^15–17^

Inserting Eq. 4 and Eq. 3 into Eq. 1, we arrive at a dynamical equation for the mean trait of a non-Gaussian trait distribution,

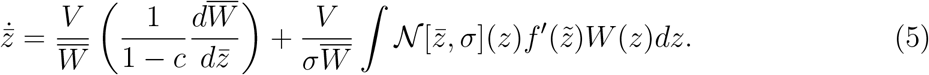

Asymptotic analysis confirms that the first term vanishes when *c =* 1 due to a first-order zero in 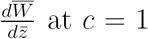 (see Appendix D). The first term in Eq. 5 corresponds to a classical fitness gradient dynamics model, and represents the complete dynamics if the trait distribution is Gaussian (*f*′ = 0).^17, 34^ The second term determines how the higher-order moments of the trait distribution affect the evolutionary dynamics. Importantly, this new term depends explicitly on the values of higher-order moments at each timestep. Therefore, Eq. 5 can only be used to determine the dynamics if additional differential equations are specified for the trait variance, skewness, etc, which makes full determination of the dynamics a “moment closure” problem because the dynamics of the *n*^*th*^ moment will, in general, depend on the (*n* + 1)^*th*^ moment.^35, 36^ However, Eq. 5 can still be used to find the mean trait equilibrium and its stability as functions of the other moments.

We next expand the trait distribution in Eq. 4 as a Gram-Charlier A series,

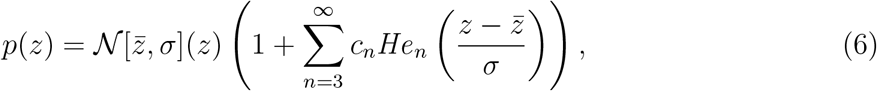

where the expansion coefficients *c*_*n*_ are uniquely determined by the moments of the trait distribution (see Appendix B). Inserting the rightmost parenthetical term into Eq. 5 as *f*, we use the property of Hermite polynomials 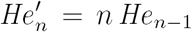 and the change of variables 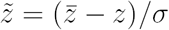 to find,

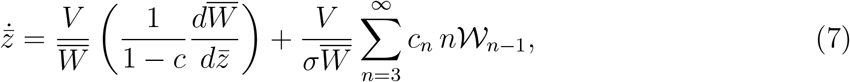

where the first-order term corresponds to classical phenotypic evolutionary theory. In the higher-order terms, the singly-indexed family of integrals 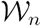 is given by

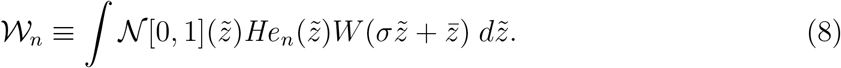

This equation represents a projection of the fitness landscape onto a basis of Hermite polynomials, with finer-scale features in the fitness landscape being represented by larger values of *n* in the series. However, if the fitness landscape is sufficiently smooth, there always exists some *n* above which the sequence 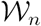 continuously decreases, suggesting that a finite set of 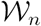 would be sufficient to describe many simple fitness landscapes.

We refer to the series 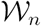 as the “coupling coefficients” because the form of Eq. 7 suggests that the values of 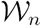 represent the degree of coupling between the fitness landscape and progressively larger cumulants of the trait distribution. Larger-scale features of the fitness landscape affect the dynamics over longer timescales, and thus appear in lower-order 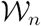; conversely, higher-order coupling coefficients provide increasingly precise information about the fitness landscape, and small-scale dynamical changes within it. Thus 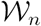 serve a similar function to the individual selection differentials described in previously-developed models of non-Gaussian breeding value distributions using multilocus genetic theory.^23, 37^

Importantly, the coupling coefficients 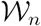 are parameters that only need to be computed once as long as the fitness landscape itself remains constant. This allows the integro-differential dynamics of an arbitrary trait distribution and fitness landscape to be expressed as a series of coupled ordinary differential equations for the time-varying moments (appearing in the individual *c*_*n*_ of the Gram-Charlier series), with the fitness landscape appearing only as a set of known coupling terms 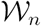. For most fitness landscapes the individual 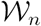 are straightforward to compute analytically (using generating functions) or numerically, due to the orthogonality properties of Hermite integrals.^38^ This allows the set of 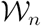 to be “pre-computed” for a given fitness landscape and then inserted as explicit terms into the dynamical equation Eq. 7. The dynamics of the trait distribution subject to this fitness landscape can then be determined without the need to perform any additional time-dependent integrals over the fitness landscape. If the fitness landscape were to change in time, any change arising from a process independent of the trait distribution (e.g. rainfall variation due to large scale weather patterns, or depletion of predators due to overfishing) would allow the time variation of each 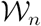 to be solved separately from the trait distribution dynamics. The resulting set of 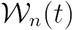 may then be inserted into Eq. 7 to yield the trait dynamics.

## 2.2 Trait variance dynamics

The method used above to derive the dynamics of the trait mean may be employed to derive corresponding dynamical equations for any moment or cumulant of the trait distribution. Here, we determine the dynamical equation for the trait standard deviation (and thus variance) using a similar method to that above.

As with the mean trait, we assume that the trait variance *M ≡ σ*^2^ has linear heritability with dynamics specified by

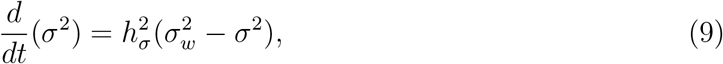

where 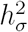 is the *variance heritability,* or the degree to which the phenotypic variance in one generation influences the phenotypic variance of the next generation. 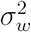 is the trait variance after selection, which is defined in a manner analogous to 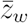. If the (unmodeled) recombination and mutation processes are sufficiently smooth, such a linear relationship and the value of 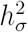 represents the lowest-order term in a suitable series expansion due to the summation properties of cumulants of random variables.^39, 40^ This relation based on early work on the infinitesimal model,^28^ in which a given trait is continuous due to an effectively infinite number of individual loci contributing to it. We further discuss below the limitations of using a linear heritability equation.

If the trait distribution has the separable form Eq. 4, then it can be shown (see Appendix E) that Eq. 9 simplifies to a dynamical equation of the form,

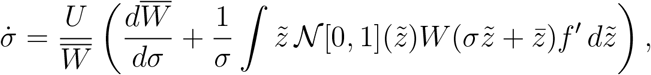

where *U* ≡ (*σ h*_*σ*_)^2^/2. Inserting the Gram-Charlier series (Eq. 6) for *p*(*z*) and exploiting the properties of the Hermite polynomials results in a final expression,

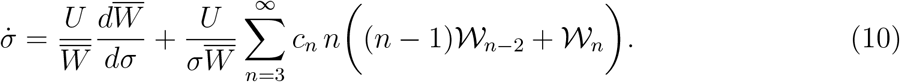

As with the mean trait dynamics Eq. 7, the variance dynamics (and thus the width of the trait distribution) simplifies to a gradient of the mean fitness, plus an infinite summation over 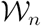 that takes into account increasingly fine-scale moments of the trait distribution. Using this construction technique, the dynamics of any arbitrary moment or cumulant of the trait distribution may be expressed as a dynamical equation with a similar form. The properties of Hermite polynomials guarantee that the fitness landscape is coupled to the dynamics of each moment solely through a fixed set of 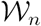, further emphasizing how the moment-based formulation abets analytical work (through truncation of the infinite series) or numerical study (through pre-computation of an arbitrary number of terms in the series of 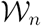).

## 2.3 Mean fitness dynamics

Based on the forms of Eq. 7 and Eq. 10, the dynamics of the mean fitness are given by

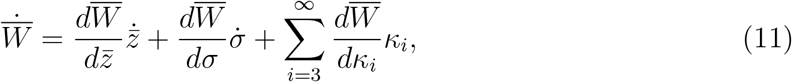

where *κ*_*i*_ are the cumulants of *p*(*z*). Because all derivatives of 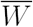 may themselves be expressed in terms of the cumulants of the trait distribution *p*(*z*), the dynamics of 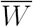 do not involve an additional dependent variable in the dynamical system. The form of 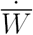 suggests that, even in the absence of additive genetic variance *(V =* 0) or in the case of a fixed mean trait 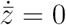, the mean fitness may continue to change due to the contribution of higher-order cumulants of the trait distribution to the dynamics. These higher-order effects are absent in the standard breeder’s equation and will discussed further below.

## 2.4 Gradient dynamics with leading-order corrections

Here, we simplify equations Eq. 7 and Eq. 10 to account for only the leading-order effects of non-Gaussian moments in the trait distribution, resulting in a closed form for the dynamical equations.

The general framework used above for deriving 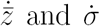 may, in principle, be used to derive dynamical equations for an arbitrary number of moments of the trait distribution. In these cases, a Gram-Charlier series with a non-Gaussian leading-kernel may be preferable.^41^ However, in the following sections we focus primarily on the dynamics of the first and second moments of the trait distribution (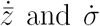 derived above) because in many standard fitness models, selection acts directly on the mean and width of the trait distribution, but not necessarily the skewness and higher moments. As a result, the terms in the series 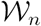 decrease quickly in magnitude, causing the dynamics of 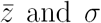 to be nearly uncoupled from the dynamics of higher moments. This is equivalent to assuming that higher-order cumulants of the trait distribution affect the dynamics solely as fixed parameter values in the series terms in Eq. 7 and Eq. 10. This restriction implies that higher moments of the trait distribution have zero effective heritability, and that natural selection, together with reproduction, mutation, and recombination, causes these moments to vary much more slowly than the mean and variance.^36, 42^ However, even if higher moments are allowed to vary continuously due to the action of natural selection, the equilibria found under our assumptions would remain the same—although their stability may be subject to additional constraints.

In order to illustrate potential applications of this formulation and to produce closed-form results, in what follows we truncate at fourth order the infinite series *c*_*n*_ appearing in Eq. 7 and Eq. 10. This fourth-order closure is used in order to isolate the effects of asymmetry (*c*_3_) and heavy-tailedness (*c*_4_) on the dynamics of the trait distribution: neither effect can be described by the original breeder’s equation due to its implicit assumption of a Gaussian trait distribution. Because of the ease with which individual 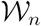 may be computed, additional terms could easily be added to account for other effects; however, the contribution of these higher-order terms to the dynamics of lower-order moments is bounded due to the scaling properties of the Gram-Charlier series (*c*_*n*_ ~ 1/*σ*^*n*^). Similar cumulant closure relations appear in moment-based models of ecological dynamics.^35, 43, 44^ In previous work by other authors,^23, 37^ the series terms in a discrete-time cumulant dynamical model were computed explicitly for the case of truncation selection, for which the short-time dynamics primarily depend on the leading moments. Additionally, in some discrete-time models of multilocus selection, the cumulant dynamical equations intrinsically contain a finite number of terms and cross terms.^45, 46^

Together, these assumptions result in a simplified set of dynamical equations,

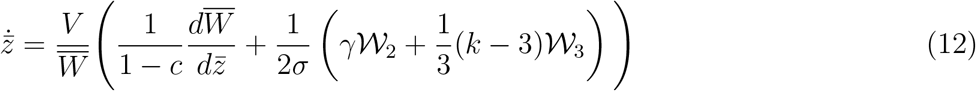

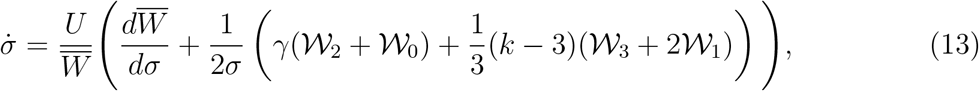

where *γ* and *k* are, respectively, the skewness and kurtosis of the trait distribution. While *γ* may take any value, the kurtosis is mathematically bounded from below by *k* ≥ *γ*^2^ + 1. Together, Eq. 12 and Eq. 13 may be considered a first-order “correction” to the classical fitness gradient dynamics equation, and they account for the leading-order effects of non-Gaussian features of the trait distribution. When the fitness landscape is centered (*c* = 1), the first terms vanish from both of these equations. In this case, if the trait distribution itself is purely Gaussian (*γ* = 0, *k =* 3), then the remaining terms vanish from the right hand sides of both equations and the trait distribution evolves under classical phenotypic evolutionary theory. However, if the trait distribution is non-Gaussian but *c* = 1, then 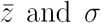 will vary due entirely to the non-Gaussian components of the trait distribution.

## 3 Assumptions and related models

Our model is comparable to moment-series approaches previously used to study natural selection in phenotypic and genotypic systems under various assumptions;^23, 36, 37, 46–48^ we review these approaches and in greater detail in Appendix L. As in previous models, we assume that the full dynamics of the trait distribution may be approximated by a finite series of ordinary differential equations, thus reducing a complex partial differential equation problem to a lower-dimensional moment evolution problem. Mathematically, the set of techniques upon which we base our analysis parallels those found in models of genetic processes under the infinitesimal model, in which a given continuous trait is assumed to depend on an arbitrary number of alleles—in particular, the use of a Gram-Charlier series as a starting point for cumulant iteration equations was pioneered in genetics by Zeng,^48^ as well as Turelli and Barton.^23^ Additionally, we note that several related works have focused on the distribution of fitness values,^46, 49^ including recent work producing the intriguing result that many fitness distributions asymptotically approach a fixed class of distributions.^36^

Our assumption of a linear heritability for higher cumulants, Eq. 9, represents the primary assumption of our model regarding the underlying mechanisms of genetic inheritance in our system; it thus introduces the primary limitations of this purely phenotypic approach because it does not include an explicit mating mechanism. For extremely strong selection (leading to large changes in the trait distribution within one generation), our model may fail due to both the continuous time assumption and the presence of higher-order, terms in Eq. 9. The form of these terms depends on the underlying genetic process, and their general form has previously been found using multilocus theory.^23, 45^ These subleading terms affect the dynamics over long timescales, and may alter the stability criteria of equilibria.

Our continuous time phenotypic equations may be compared to previous work on cumu-lant dynamics that treat selection as a discrete-time repeated sampling process weighted by the fitness.^50^ Thus we test our findings below against a simulated Wright-Fisher process, and we find general agreement (see Appendix N for details of this numerical work). We thus emphasize that our model is best applied to the study of short-term phenotypic evolutionary trends when genetic assays are unavailable or infeasible; however, over longer timescales in which the additive genetic variance parameter *V* varies, we expect high-order effects in her-itability to manifest. These limitations are consistent with the classical usages of “gradient dynamics” models (which our model essentially generalizes), which have found particular utility for the study of coupled ecological and evolutionary processes.^15–17, 34^

## 4 Results

### 4.1 Cryptic forces of selection arise from non-Gaussian trait distributions

In classical phenotypic evolution, the first term in Eq. 12 is associated with the “force of selection” on the mean trait.^51^ However, the remaining terms in Eq. 12 and Eq. 13 show that the trait distribution can change even when this term is zero, allowing the trait distribution to evolve in the absence of any apparent force of selection. These cryptic selection forces can have significant effects on the overall dynamics of natural selection.

As a demonstration of this effect, Figure 1 shows the behavior of the system Eq. 12, Eq. 13 relative to a null model in which the trait distribution is always Gaussian (in which case all terms containing 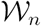 vanish in the dynamical equations). The figure illustrates two separate cryptic forces of selection: the excess selection force on the mean trait (left plots) and the excess selection force on the trait variance (right plots). For the Figure, we use a simple fitness landscape consisting of exponential directional selection,^40, 52, 53^

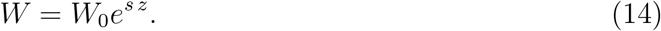

Such a landscape represents limiting case in several common contexts, including selection on metric traits (which frequently have logarithmic ranges),^18, 21^ evolution of biochemical reactions subject to microscale energetic constraints,^54, 55^ and cases in which fitness scales with mutation count.^56, 57^The parameter *s* in Eq. 14 describes the relative strength of selection on the trait *z.* For this fitness landscape, the individual coupling coefficients are observed to obey the general relationship 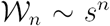 (Appendix H); thus the broadest features of the fitness landscape (lowest order 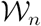) contribute the most to directional selection.

**Figure 1.**
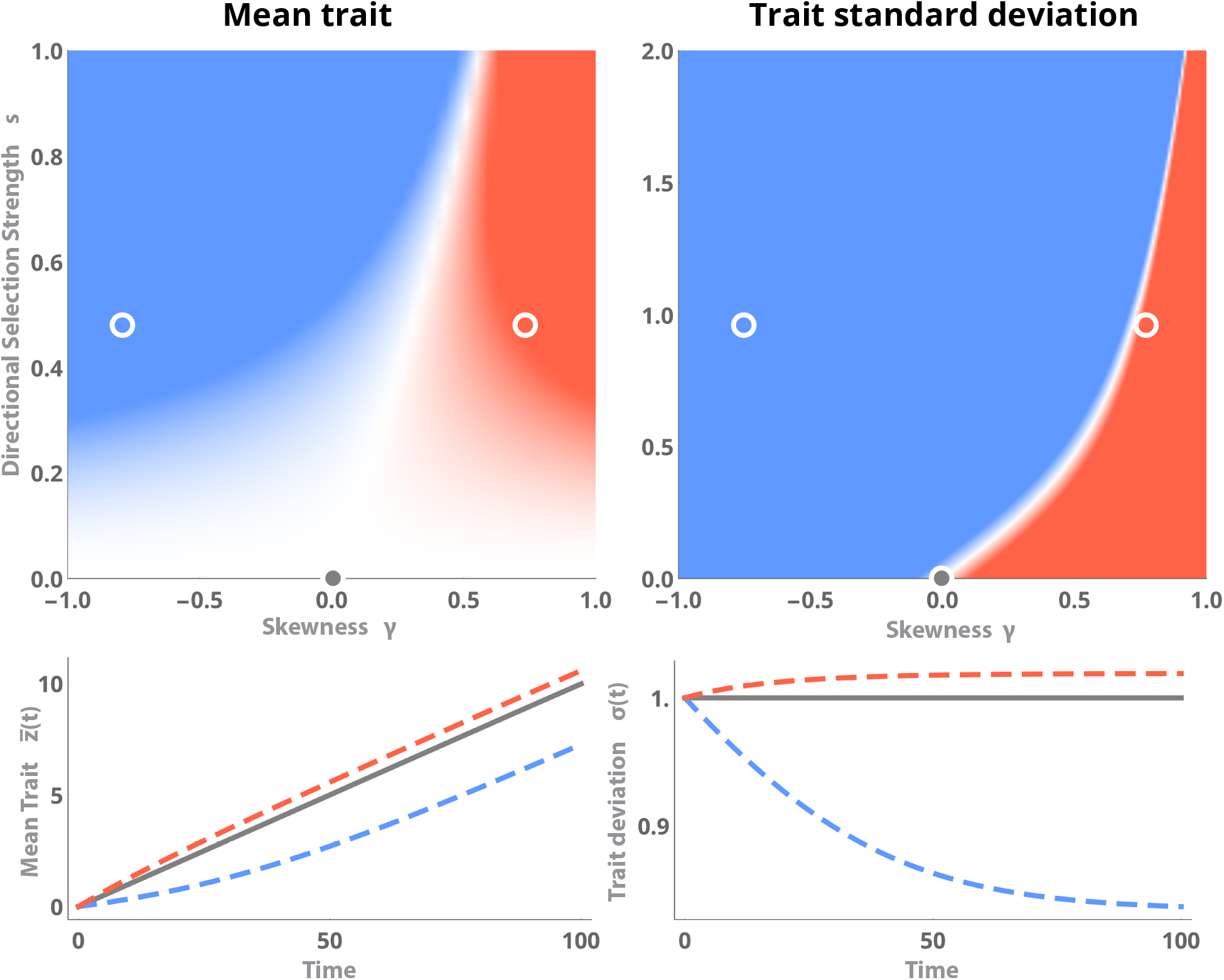
Cryptic forces of selection under directional selection. Top: The excess force of selection due to the non-Gaussian form of the trait distribution, for the mean trait dynamics (left) and trait standard deviation dynamics (right). Colored shading indicates the relative direction and magnitude of the cryptic terms in Eq. 12 and Eq. 13, with red (blue) indicating cryptic forces that accelerate (retard) the growth of each moment. Lower plots represent example dynamics of the mean and standard deviation for representative points on each color plot. White circles on upper plots indicate parameter values (*γ*, *s*) used for the trajectories in the lower plots: the grey trace indicates classical fitness gradient dynamics with an unperturbed Gaussian trait distribution, while red and blue traces indicate cases in which higher trait moments speed or hinder the evolutionary dynamics, respectively.

The upper portion of the figure describes the relative magnitude of the excess force of selection as a function of the trait skewness γ and the selection strength *s*; for simplicity, the kurtosis *k* is held fixed at its theoretical minimum *k* = γ^2^ + 1. At the origin, the trait distribution is Gaussian and so the cryptic selection force is zero; however, as either the skewness γ or the selection strength *s* increase, the relative contributions of the non-Gaussian terms in Eq. 12, Eq. 13 increases. Red regions correspond to cases in which the cryptic selection force is positive (and thus assists directional selection), whereas blue regions correspond to cases in which the non-Gaussian contributions retard directional selection. The plot suggests that positive skewness (corresponding to a trait distribution with a long tail of large trait values) generally speeds the evolutionary dynamics of both the trait mean and trait variance, primarily due to the added contributions of extremal individuals. The opposite is true for negatively skewed populations in which most individual traits exceed the trait mean, due to outlier individuals producing offspring with lower fitness.

In the lower portions of the figure, trajectories of 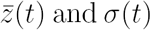 are shown for a numerically-constructed non-Gaussian trait distribution (Appendix M) using (γ, *s*) values marked by open circles in the blue and red regions of the upper plots; for comparison, a trajectory consisting of the “null” case of a Gaussian trait distribution is shown in gray. The dynamics of each distribution relative to the Gaussian case proceed as predicted by the magnitude of the cryptic force, with the primary advantage/detriment due to skewness occurring initially before the rate of evolution eventually stabilizes. Notably, directional selection causes a continued increase in the mean trait 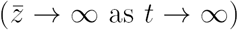, but the trait distribution width *σ* stabilizes to a constant value that is proportional to the skewness. Thus the effect of non-Gaussian features in the trait distribution may manifest experimentally as a constant variance that is larger or smaller than that predicted under classical evolutionary theory.

In the Appendix, we compare these results to Wright-Fisher simulations of phenotypic evolution in a population initialized to the same starting values of the mean, variance, skewness, and kurtosis as was used in these equations, and we find general agreement. In the simulations, as in real populations, the values of the skewness and kurtosis drift over time due to accumulated sampling errors (this corresponds to a timescale over which the “fixed cumulant” assumption above is no longer valid). However we observe that trait means and variances tend to drift monotonically under exponential directional selection, and so even over long timescales the calculated direction of the cryptic force of selection correctly predicts the dynamics relative to the Gaussian distribution.

### 4.2 Transient evolutionary responses to stabilizing and disruptive selection

In addition to generating qualitatively different dynamics, non-Gaussian features of the trait distribution may affect the long-term duration and direction of natural selection. In order to illustrate this effect in a fitness landscape with a variable number of local maxima, we next consider a general fitness landscape described by an arbitrary polynomial,

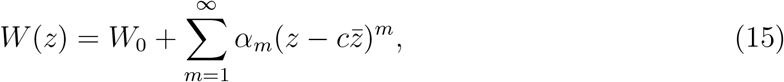

where the relative magnitudes of the various coefficients *α*_*m*_ determine the number of local maxima and minima of the landscape. In general, if the largest non-zero *α*_*m*_ is positive, then for large values the fitness landscape looks like a 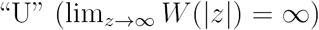, and so the value of the mean trait 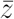 eventually diverges: natural selection proceeds indefinitely in the system as the mean trait continuously increases. Sub-leading terms in the polynomial Eq. 15 induce short term transients that may affect the dynamics only temporarily depending on the initial conditions. Globally, however, natural selection will proceed continuously in a U-shaped landscape, as has been reported experimentally.^58^ Conversely, if the largest non-zero *α*_*m*_ is negative, then the fitness landscape looks like a hill at large *z* and so the the mean trait and trait variance will always eventually equilibrate at an intermediate stable solution—in which case natural selection proceeds transiently until this solution is reached.^59^

We can investigate the manner in which skewness and kurtosis affect the timescale of natural selection by studying the specific case of a quartic fitness landscape under transient natural selection (*α_m_ =* 0 for *m* > 4; *α_4_ <* 0). This landscape can have either one or two local maxima depending on the relative magnitudes of *α*_2_ and *α*_4_ in Eq. 15. Thus for a continuous, unbounded trait, a quartic fitness landscape represents the simplest landscape that can model both stabilizing (one maxima) or disruptive (two maxima) selection.^34^

For each value of the skewness *γ* and kurtosis *k*, the equilibrium solutions of Eq. 12 and Eq. 13 are independent of the initial conditions and can be found analytically along with the Jacobian matrix describing their local stability. Figure 2 shows plots of the maximum eigenvalue of this Jacobian matrix associated with the intermediate phenotype, parametrized by the skewness *γ* and kurtosis *k* (the kurtosis lower bound *k > γ*^2^ + 1 is indicated by a solid gray line). For cases in which the dynamical equations yield multiple solutions, the intermediate equilibrium phenotype is defined as the solution of Eq. 12 and Eq. 13 with 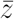 closest to 0. In the blue regions on the plots, this intermediate phenotype is stable and so the mean trait approaches this point. In the red regions of the plots, the intermediate phenotype is unstable and so the mean trait instead approaches another, extremal equilibrium point. The relative darkness of the plot colors represents the relative speed of the evolutionary dynamics: darker blue indicates that the intermediate phenotype is achieved relatively quickly, while darker red indicates that extremal phenotypes are reached quickly.

**Figure 2.**
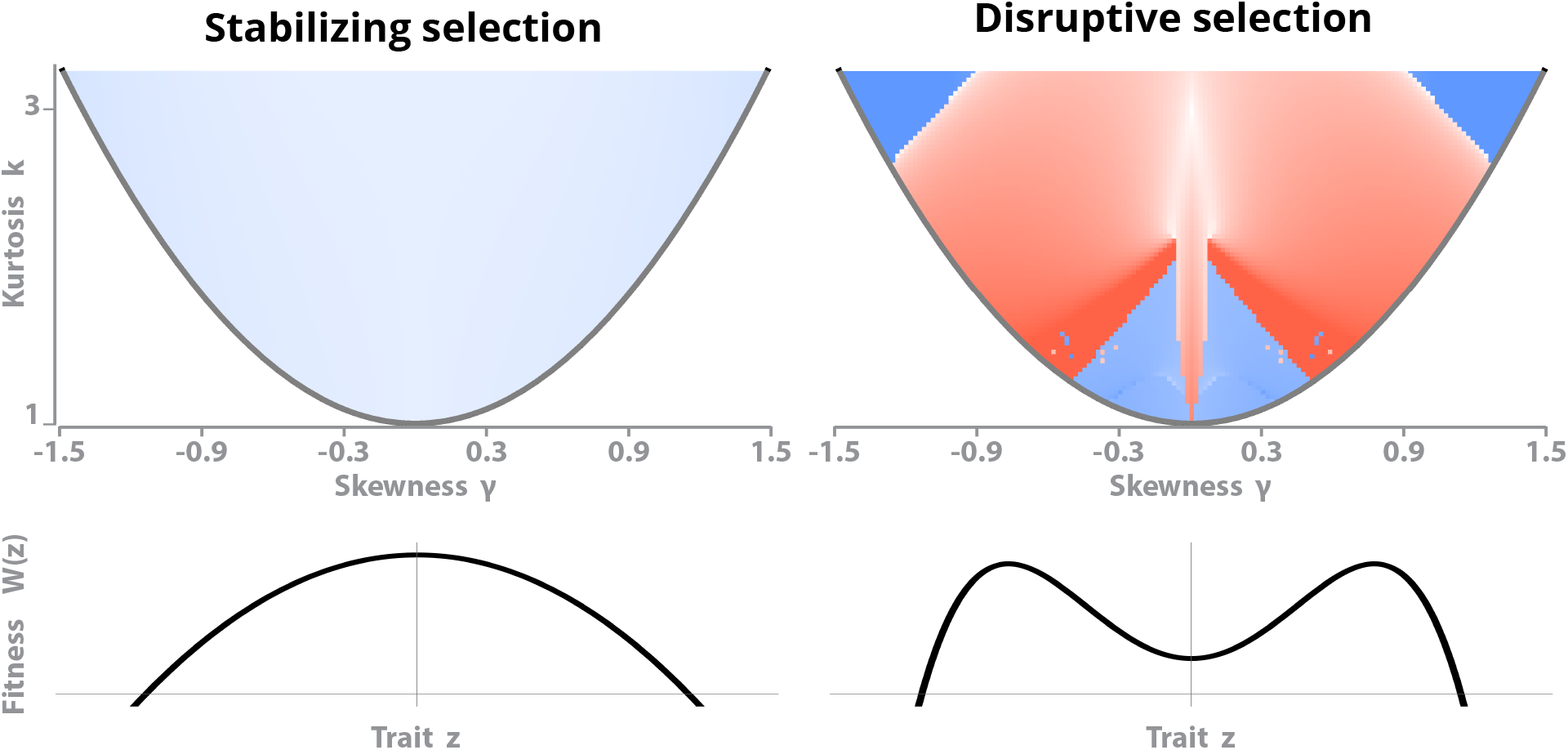
Evolutionary dynamics under stabilizing or disruptive selection. Colored shading represents the maximum eigenvalue associated with the Jacobian matrix of 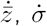 on a logarithmic scale, evaluated at the equilibrium nearest to 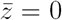 (the intermediate phenotype) and parametrized by the skewness and kurtosis of the trait distribution (all other parameters are held constant). Negative maximum eigenvalues (blue regions) represent dynamics that eventually converge to the intermediate phenotype, positive maximum eigenvalues (red regions) represent dynamics that approach an extremal phenotype, and the intensity of shading indicates the instantaneous rate of the dynamics. The solid gray line indicates the lowest mathematically-valid value for the kurtosis, *k > γ*^2^ + 1. Beneath each figure is a diagram of the fitness landscape *W*(*z*) that produced it. For this figure, *c* = 1, *V =* 1, *U =* 10 in Eq. 12 and Eq. 13. Shading ranges from eigenvalue values of *−*2 (dark blue) to 2 (dark red).

The points *γ* = 0,*k* = 3 on each plot correspond to the default case of a purely Gaussian trait distribution with a time-varying mean and standard deviation. For stabilizing selection this point resides within a blue region, consistent which the intuitive result that a Gaussian trait distribution evolves towards the location of the global maximum of the fitness landscape, and that the range of trait values decreases over time as more and more individuals in the population approach this optimum (*σ*(*t*) → 0).^54^ Indeed, for stabilizing selection, non-Gaussian features in the trait distribution only barely affect the rate of the dynamics, consistent with experimental results suggesting that traits under stabilizing selection universally attain intermediate optima.^60^ Additionally, this suggests that if *γ* and *k* were themselves dynamical variables that changed either due to selection or mating effects, the mean fitness would nonetheless always reach a finite value dictated by the maximum of the fitness distribution.

In a disruptive landscape, the intermediate phenotype is disfavored for almost all trait distributions near the classical case of a Gaussian distribution at *γ* = 0, *k =* 3 (red regions in Figure 2). This is expected because the maxima of the disruptive landscape occur away from the intermediate phenotype at *z =* 0, and so the trait moves towards these dispersed values in the absence of additional destabilizing dynamics. However, when the trait distribution is strongly non-Gaussian, the intermediate phenotype near *z =* 0 regains stability, an effect that is particularly pronounced when the trait distribution is strongly asymmetric and flat (large *γ*, *k* ≈ *γ*^2^ + 1). High skewness and kurtosis correspond to the trait distribution containing a relatively large fraction of individuals with extremal phenotypes, which represent a large enough trait range that the distance between the two symmetric equilibria in the disrupting fitness landscape becomes relatively insignificant (the locations of these off-center equilibria depend on *α*_2_/*α*_4_ in Eq. 15). As a result, the overall average phenotype returns to the center due to the leading-order effect of the negative *α*_4_ term in Eq. 15.

In the Appendix, we compare the dynamics predicted by local stability analysis of Eq. 12 and Eq. 13 with the dynamics of a population subject to Wright-Fisher dynamics with matching starting cumulants as used here. We find that the short-time dynamics of the Wright-Fisher process are directly analogous to those predicted by the local analysis, particularly because additional effects (such as variation in the genetic variance *V*, and fluctuations in the values of higher cumulants due to sampling drift) do not manifest over the infinitesimally-short timescales which local stability analysis applies.

Significant skewness and flatness in the trait distribution can obscure the effects of disruptive selection and potentially serve as a mechanism to preserve or enhance phenotypic variance, even when selection itself acts on the trait variance—potentially implicating higher trait moments in speciation and ecological phenomena that depend on the phenotypic vari-ance.^19^ Although the dynamical equations would become substantially more complex if the skewness and kurtosis also responded to selection, results from other cumulant dynamical systems^39, 40^ suggest that genetic or mating processes that preserve an arbitrarily high order moment would result in similar “freezing” of the dynamics of lower-order cumulants. Thus, even if natural selection alters arbitrarily high moments of the trait distribution, if mutation or mating serve as a “source” that constrains an arbitrary moment of the trait distribution, then lower-order moments such as the phenotypic variance would also be prevented from reaching zero in certain regions of parameter space. This process represents a generalization of the concept of a “mutation-selection balance,” a concept typically invoked in order to justify holding the phenotypic variance fixed during selection.^15, 36, 46^ However, as with the traditional breeder’s equation, an additional equation specifically incorporating the underlying genetics would be required in order to justify this process biologically.^47^

### 4.3 Time-variation of the heritability

Experimental studies of phenotypic evolution in artificial selection regimes generally quantify genetic effects using the narrow sense heritability, *h*(*t*)^2^ ≡ *V/σ*(*t*)^2^, defined as the ratio of additive genetic variance *V* to overall phenotypic variance *σ*^2^. In large populations, *h*^2^ changes slowly enough that it can be estimated without the need for explicit identification of a trait’s genetic origin. However, recent experimental results have suggested that *h*^2^ may change appreciably during short periods of strong selection, especially when the underlying genetics (and thus *V*) exhibit complex dynamics.^21^ In particular, rapid changes in the phenotypic variance *σ*^2^ may underly this phenomenon over short timescales,^19, 60^ even when an insufficient number of generations have elapsed for *V* to change appreciably.

We note that by including changes in higher moments than the mean, our framework naturally describes changes in the observed heritability arising from the dynamics of trait variation. In order to study this effect, we re-parametrize our model in terms of quantities more readily measured experimentally: our two dynamical variables 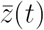 and *σ(t)* may be replaced by the equivalent conjugate variables of heritability *h^2^(t)* and mean fitness 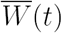 (see Appendix G). Because the exact underlying trait values or distributions may not be accessible in certain experimental contexts, these variables are more descriptive of macroscopic population trends in population-level assays, such as long-term bacterial evolution experiments.^61^

In Figure 3, we show the short-time evolutionary dynamics of the heritability under a linear directional fitness landscape,

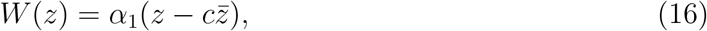

for which the coupling coefficients are computed in Appendix K. Because any arbitrary, smooth fitness landscape may be approximated as linear in the neighborhood of the mean trait, this parametrization illustrates a general relationship between the heritability and mean fitness dynamics over short timescales. Setting *α*_1_ > 0 results in positive directional selection 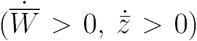 because regions of a fitness landscape with positive slope tend to drive a population towards a higher mean fitness. The solutions to Eq. 12, Eq. 13 corresponding to each timepoint are shown in Figure 3A, where they are parametrized in terms of the heritability as a function of mean fitness (which itself increases in time). In the figure, different traces correspond to various values of the skewness *γ* and kurtosis *k.*

**Figure 3.**
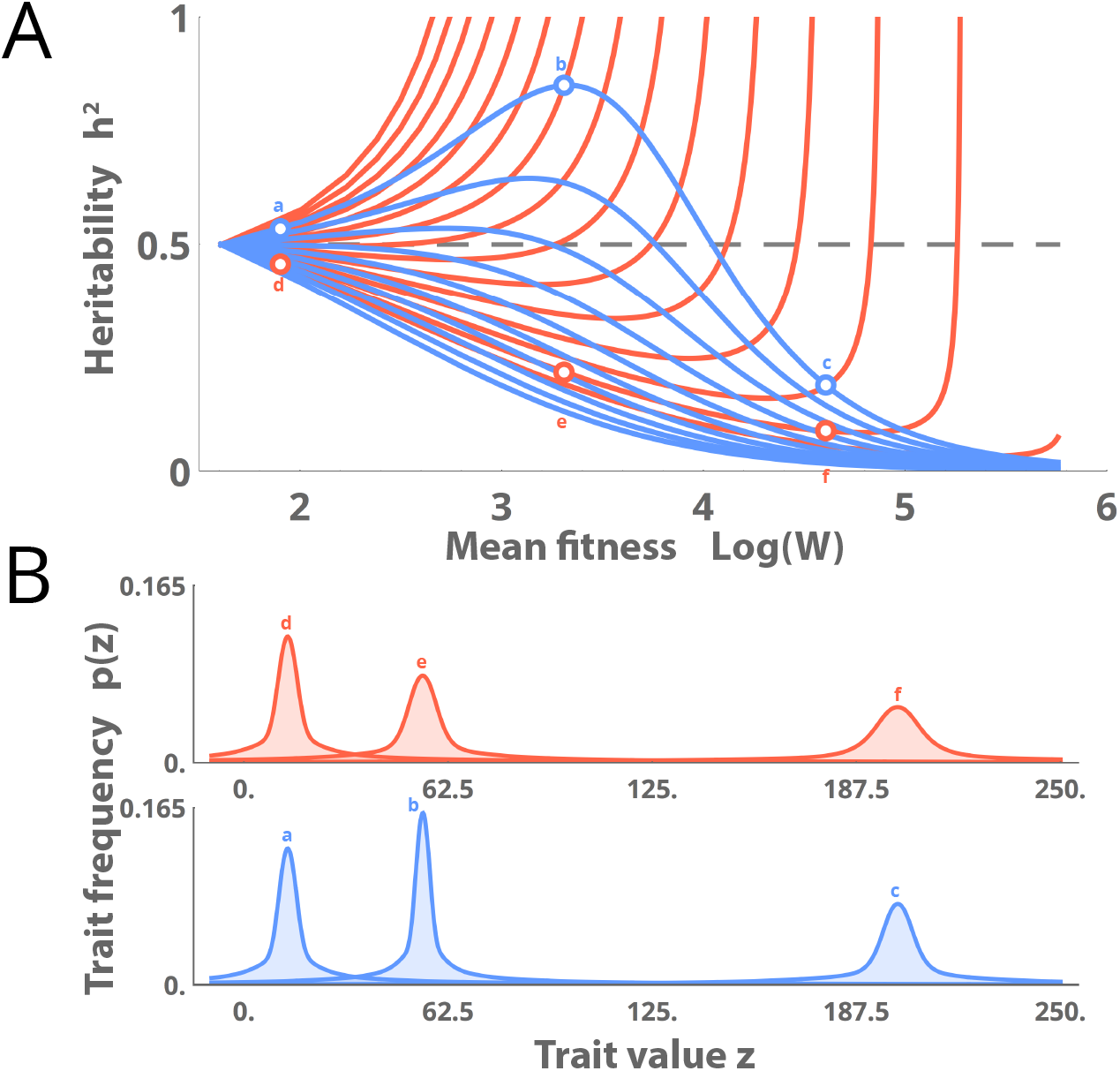
Time-dependent heritability under directional selection. (A) The heritability *h*^2^ as a function of the mean fitness, which increases continuously over time in a directional fitness landscape. Colors represent cases in which the trait distribution skewness is *γ* = –0.05 (red) or *γ* = 0.05 (blue). Different traces of the same color correspond to increasing values of the kurtosis *k* in the range *k = γ*^2^ + 1 to *k = γ*^2^ + 6, with slower timescales (lower traces) corresponding to larger values of the kurtosis. The gray dashed line corresponds to evolutionary dynamics with constant heritability, which the dynamical equations recreate when *γ* = 0, *k =* 3. (B) The trait distributions for three representative timepoints (and thus values of 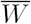) marked by open circles in (A). For this figure, *α = V =* 1, *U =* 0.01, *c* = 0.5.

In general, a Gaussian trait distribution (*γ* = 0, *k* = 3 in Eq. 12 and Eq. 13) always produces the default case of constant heritability (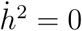, gray dashed line in Figure 3A). An evolving population with positive skewness (blue traces) exhibits heritability that eventually decreases in time, primarily due to a high fraction of outlier individuals in the high-fitness tail of the trait distribution. These individuals produce higher-fitness offspring quickly enough that the overall trait range increases in time, leading to a corresponding decrease in the overall heritability—which is consistent with several field studies showing that increasing mean fitness also increases trait variation and lowers the observed heritability.^60, 62^ This growth of the variance due to tail effects is consistent with prior analytic solutions for the case of directional selection;^37, 63^ however we note that we hold the skewness and higher moments fixed in these calculations. Intriguingly, we find that high values of kurtosis retard this process at short timescales, producing some scenarios where the heritability appears to increase transiently, before eventually relaxing non-monotonically to zero over longer timescales (topmost blue trace). Such non-monotonicity in the heritability under directional selection would be an observable experimental signature of non-Gaussian dynamics.

Conversely, a population with negative skewness (red traces in Figure 3A) has a long tail of individuals with comparatively low fitness; these individuals serve to counteract the tendency of directional selection to increase trait ranges, and thus cause the trait variation to decrease in time—leading to an accompanying increase in the trait heritability. Because this effect is driven by the lower-fitness tail of the distribution, it may partly explain experimental results that have reported a disproportionately large contribution of rare phenotypes to the heritability of certain deleterious traits.^64, 65^ The differential effect of left and right skewness is further apparent in the trait distributions at three representative timepoints in the dynamics shown in Figure 3B, where the apparent width of the trait distribution varies non-monotonically and causes continuous changes in the heritability observed in the upper portion of the figure. This substantial variation in the distribution’s width at various points in the dynamics may confound efforts to study experimentally the evolution of ecological niche width,^66^ as non-Gaussian features in the trait distribution may cause transient contraction and expansion of the observed trait range, even in the absence of competition.

In the Appendix, we perform a comparable set of simulations using Wright-Fisher dynamics, and we find general agreement in the dynamics, including non-monotonicity in the dynamics at large values of *k,* as well as a qualitative shift from *h*^2^ → 1 to *h*^2^ → 0 when *γ* changes sign from negative to positive.

## 5 Discussion

We have presented a formulation of phenotypic evolution that seeks to iteratively relate the dynamics of an arbitrary trait distribution to an arbitrary fitness landscape. This simplified model explains several phenomena observed in numerical simulations of non-Gaussian trait distributions evolving in simple fitness landscapes. We have shown that skewness or asymmetry in the trait distribution can delay directional selection or prevent disruptive selection, and that the heritability—typically assumed to be constant over short timescales—can vary in time with complex, non-monotonic dynamics as the mean fitness changes.

Our results are consistent with empirical and theoretical analyses that formulate phenotypic evolution in terms of the probability distribution of fitness values *p*(*W*) rather than in terms of an explicit trait distribution *p*(*z*) and auxiliary fitness function *W*(*z*)^49^ However, the trait-based model here is more readily applied to arbitrarily complex fitness functions (particularly non-invertible and multimodal fitness landscapes),^46^ and thus may assist the study of explicitly experimental quantities such as selection coefficients and realized heritability. For fitness distribution models, similar cumulant dynamical equations have been derived under distinct constraints and approximations, and for certain types of experiments (e.g., microbiological lineage assays) these formalisms may be preferable.^36, 40, 46, 61^

Our explicitly phenotypic approach does not involve an explicit underlying genetic model; we assume a linear heritability relation and note that this approximation holds over the relatively short timescales and slow selection regimes observed in our simulations.^28^ A more detailed model would include explicit information about the mechanics of inheritance, and how these parameters contribute to the breeder’s equation and determine the parameters it contains. An important starting point for such work would be models of non-Gaussian evolution based on a multilocus genetic models,^37, 47^ which have shown that the dynamics of the phenotypic moments may vary appreciably depending on mating effects, transmission effects, and whether selection occurs before or after transmission^23^

Other potential improvements include alternative series closure schemes to raw truncation that depend on the type of fitness landscape being evaluated, as well as additional cross terms in the dynamical equations that account for assortative mating effects. Additionally, we have assumed that the additive genetic variance remains fixed during the phenotypic dynamics; however over long timescales natural selection and mating may affect this value considerably. Coupling genotypes and phenotypes may also require the phenotypic portion of the model to be generalized for multivariate traits by defining a fully time-dependent phenotypic variance-covariance matrix, which may be particularly important for selection experiments in which the traits with the strongest selection responses are unknown a priori.^13, 27^

Another limitation of our approach comes from the truncation approximations necessary to make the dynamical equations closed-form. Here, we have chosen canonical directional and minimal stabilizing/disruptive fitness landscapes, in order to illustrate the simplest non-trivial dynamics arising from our model and to justify our inclusion of only the skewness and kurtosis (which represent the leading-order departures from Gaussian trait distributions). Additionally, we have retained only two of the dynamical equations in our model: one for the mean trait and one for the trait variance. But for certain types of initial trait distribution or fitness landscape, the leading order equations Eq. 12 and Eq. 13 may not be sufficiently accurate, and instead the full dynamical equations Eq. 7, Eq. 10, and their equivalents for higher cumulants may be necessary. The number of necessary dynamical equations, and the number of terms to include in them, depends primarily on the form of the coupling coefficients 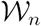, which may be found as long as the fitness landscape is known. A more detailed fitness landscape, such as one derived empirically, may require retention of more terms in the series 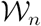, because higher-order terms correspond to finer-scale fitness landscape variations which affect dynamics over smaller length and timescales. However, we note that because computing 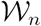 simply requires projecting a known fitness landscape onto a polynomial basis, these coupling coefficients may easily be found numerically if the parameter values for the initial trait distribution and fitness landscape are known.

## 6 Acknowledgements

W. G. thanks the NDSEG Fellowship program for support.

## 8 Appendix

## A Supplementary Figures

## B Dynamical equations for cumulants and moments

Standard derivations based on non-overlapping generations (discrete time) start with the breeder’s equation, which supposes that the mean trait distribution in the next generation depends linearly on the effect of selection on the current generation,

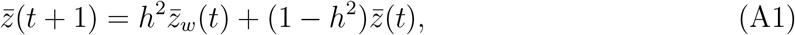

where the linear parameter *h*^2^ represents the narrow-sense heritability of the trait *z.* Rearranging the terms in this equation and taking the limit of vanishingly small generation time leads to the dynamical equation,

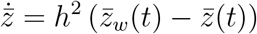

where the mean reproductive rate has been used to make time dimensionless.

One way of deriving Eq. A1 involves expressing the full distribution *p*(*z*, *t* + 1) as an integral over the distribution at the previous timestep, *p*(*z*, *t*). Various “survival kernels” and “mating kernels” are included within this integral, and they determine the contribution of mating preference, fitness, and even migration to the form of the trait distribution.^67, 68^ In these models, the linear form of Eq. A1 originates from the convolution of multiple probability distributions, which in general will result in a new distribution with a mean given by a linear combination of the means of the original distributions.

In general, the convolution of two arbitrary distributions has a distribution with cumulants given by the sum of the cumulants of the original variables’ distributions. This property is responsible for the linearity of Eq. A1, and its importance for the generation of cumulant dynamical equations for evolution has been noted by Burger.^37^ For this reason, the analogy to Eq. A1 for higher-order cumulants is given by,

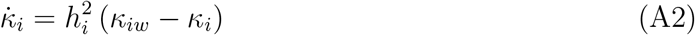

where 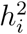 represents a generalization of the heritability for higher-order cumulants (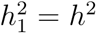 is the standard narrow-sense heritability). This equation represents an extension of the implicit assumption in the breeder’s equation that the trait response depends linearly on the strength of selection: if the means respond linearly, then other cumulants must as well as long as the trait has a well-defined probability distribution in the current generation. Linear heritability equations for the variance in the form of Eq. A2 have previously been derived explicitly for the “infinitesimal” genetic model (in which a given continuous trait arises from the independent contributions of infinitely many distinct loci) by Bulmer.^28^ Similar relations of this form are found by Turelli and Barton in both few-locus and infinitesimal genetic models (See Eq. 45 of Turelli and Barton 1994,^23^ and Eq. 3.2a,b of Turelli and Barton 199040).

While the cumulants of a random variable have the advantage of being additive, the moments of a probability distribution often provide a more familiar description of the variable. Fortunately, the dynamical equations Eq. A2 may readily be converted into dynamical equations for the central moments (*µ*_*i*_) using standard reference formulas; we note however that the first three cumulants are directly proportional to the first three central moments:

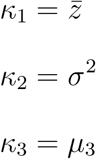

The fourth and higher cumulants have more complicated expressions given by Bell polynomials, *B*_*n*,*k*_, of the various lower-order central moments,

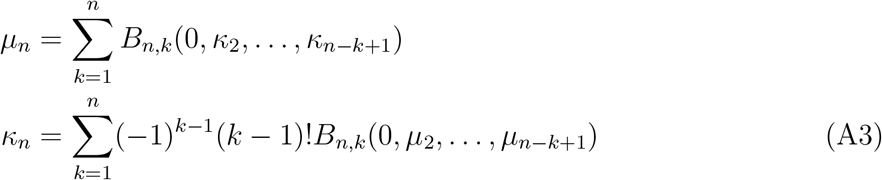

These equations may be inserted directly into Eq. A2 to obtain dynamical equations in terms of central moments. Dynamical closure in this context thus represents an assumption that cumulants above a certain order are not heritable: 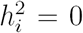 for some *i* > 1. This results in the infinite tower of coupled moment equations given by Eq. A2 becoming finite, because the explicit form of the *n*^*th*^ cumulant only depends on the *n*^*th*^ and smaller central moments.

The heritability requirements above suggest how each cumulant of an arbitrary trait distribution may have its own dynamical equation. Importantly, each of these dynamical equations may itself depend on an arbitrary number of cumulants of the trait distribution, depending on the current form of the distribution. This is because an arbitrary trait distribution may be expressed in terms of a Gram-Charlier A series,

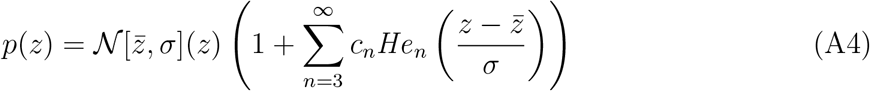

where each *c*_*n*_ is defined explicitly in terms of the moments or cumulants of *p*(*z*),

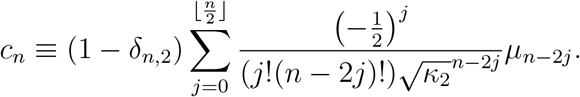

Here *δ*_*n*,*m*_ is the Kronecker delta function (*δ*_*n*,*m*_ = 1 when *n = m*; *δ_n,m_ =* 0 otherwise) and the central moments *µ*_*j*_ are given in terms of the cumulants by Eq. A3.

## C Simplification of trait dynamics when 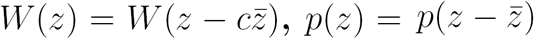

The mean fitness is given by

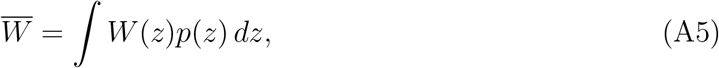

Taking the derivative of both sides of this equation,

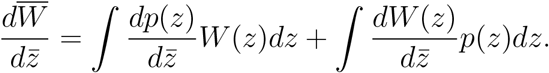

We assume that 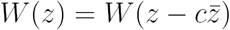

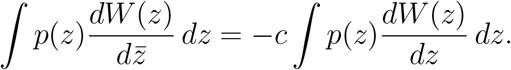

Performing integration by parts,

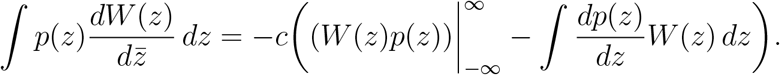

The first term on the left hand side vanishes due the compactness of *p*(*z*),

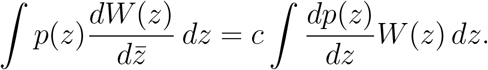

If 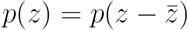 then,

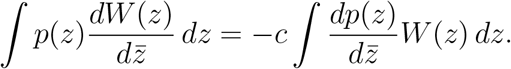

Inserting this expression into Eq. A5 results in a final equation,

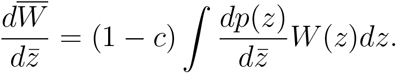

This result is intuitive for *c* = 1: if the fitness landscape depends only on the difference between each individual trait the mean trait, then the mean fitness itself cannot depend on the mean trait.

## D Behavior of 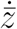 as *c →* 1

We start with the definition of the mean fitness,

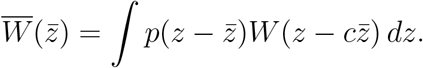

Taking a series expansion of this equation around *c* = 1,

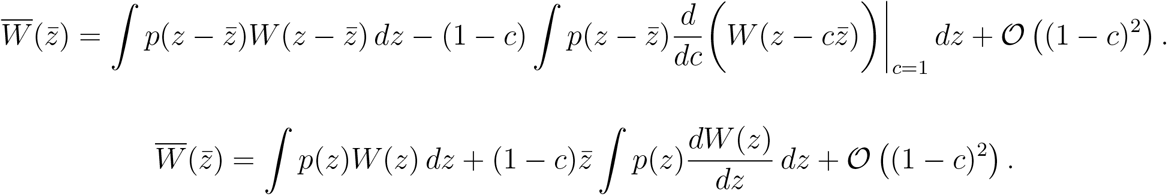

We take the derivative of both sides with respect to 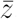,

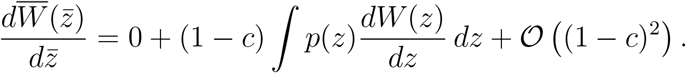

Next we multiply both sides by 1*/*(1 *- c*) and take the limit *c →* 1. We note that all terms 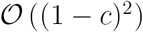 vanish in this limit,

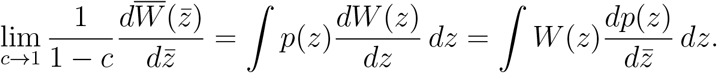

Importantly, this suggests that the dynamics of 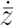 do not diverge as 1*/*(1 *- c*), because 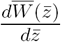 has a leading-order zero, namely (1 *- c*).

Rearranging the asymptotic equation results in the dynamics when *c =* 1,

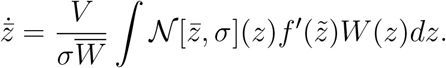

The behavior of the series in the vicinity of *c* = 1 can be found by including a higher-order term in the Taylor series found above,

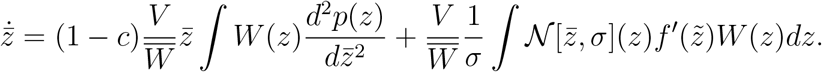

## E Derivation of trait variance dynamics 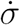

As with the trait mean, we assume that the trait variance *M ≡ σ*^2^ has linear heritability with a dynamical equation given by

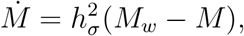

where 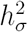 is the *variance heritability*; it determines the degree to which the phenotypic variance in one generation influences the phenotypic variance of the next generation. We expand this equation as

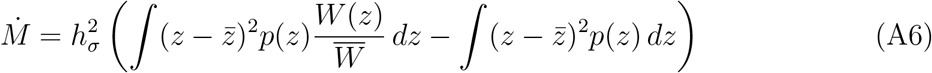

We use the substitution,

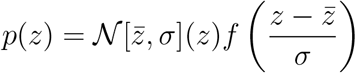

Taking the derivative of both sides of this expression results in the relationship

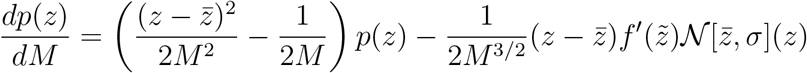

Rearranging this equation results in the expression

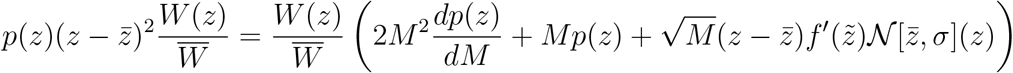

Substituting the right hand side of this expression into the integral in Eq. A6 results in the equation,

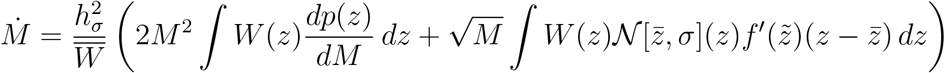

Using the substitution *M* = *σ^2^, dM* = 2*σ dσ* results in a dynamical equation for the standard deviation of the trait distribution,

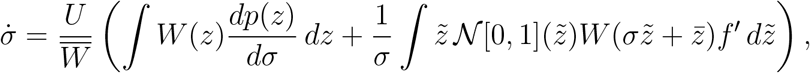

where *U* ≡ (*σh*_*σ*_)^2^/2. We note that in many standard real-world contexts and theoretical frameworks, the fitness *W* does not explicitly depend on *σ,* and so *dW/dσ =* 0, resulting in a simpler form for this dynamical equation,

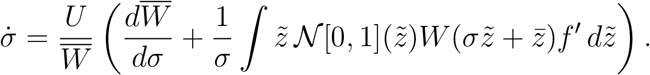

Inserting the Gram-Charlier series (Eq. 6) for *p*(*z*) results in an infinite series of the form

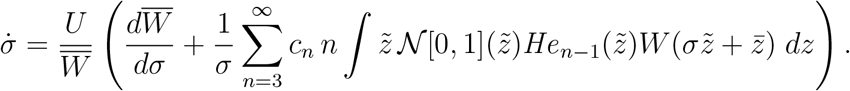

Using the properties of Hermite polynomials, this equation simplifies to

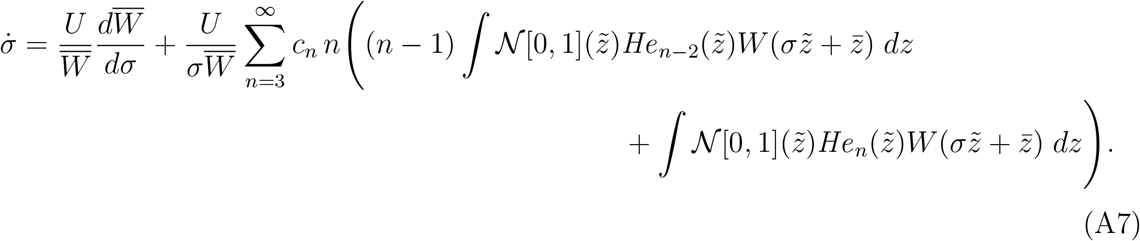

## F Dynamics of mean fitness

As the trait distribution *p*(*z*) changes in time, the mean fitness 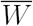 also changes. These dynamics may be expressed in terms of the trait distribution cumulants or moments by noting that the fitness landscape *W*(*z*) may be expanded in terms of an infinite series around *z* = 0,

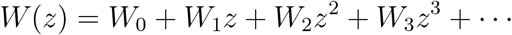

Thus the mean fitness has the form

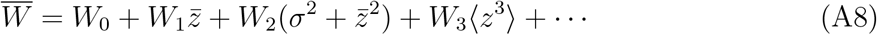

where the relationship 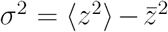 has been used. The remaining individual raw moments 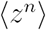 may be expressed in terms of the variance *σ*^2^ and remaining cumulants *κ*_*n*_ using the relation Eq. A3.

For a given fitness landscape, the individual cumulants (which may vary in time) may be substituted into the expression for 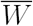 given by Eq. A8. The resulting explicit expression for 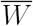 may then by substituted into the cumulant dynamical equations in order to express the dynamics in a form where the cumulants are the only dynamical variables. Depending the fitness landscape being used, the resulting equations may have a particularly simple form.

While the number of time-dependent dynamical variables required to study the trait distribution is simply the number of moments or cumulants that vary in time, for some fitness landscapes the dynamics become more tractable if 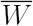 is instead treated as its own dynamical variable, with dynamics given its total derivative with respect to the remaining dynamical variables,

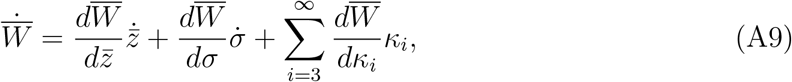

where *κ*_*i*_ are the cumulants of *p*(*z*). Because all derivatives of 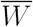 may themselves be expressed in terms of the cumulants of the trait distribution *p*(*z*), the dynamics of 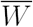 do not represent an additional dependent variable in the dynamical system. This form of 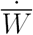 suggests that, even in the absence of additive genetic variance (*V* = 0) or in case of a fixed mean trait 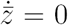, the mean fitness may continue to change due to higher-order cumulants of the trait distribution contributing to the dynamics. These higher-order effects are absent in the standard breeder’s equation and will discussed further below.

Finally, we note that under the specific closure assumptions described above and in the main text, the cumulants series *κ*_*n*_ and thus the Gram-Charlier coefficients of the trait distribution *c*_*n*_ are assumed to truncate at finite order, and the only dynamical moments of the trait distribution are assumed to be the mean 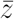 and width/deviation *σ.* The latter assumption either entails that higher moments of the trait distribution are not heritable due to specific features of recombination or mating dynamics (which are not explicitly modelled here), or—less restrictively—that the fitness landscape does not affect these higher moments. This allows Eq. A9 to have a closed form due to truncation of the infinite series,

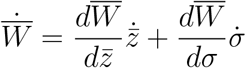

For fitness landscapes in which the dynamics of 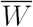 cannot be solved explicitly, this expression may be used as an additional dynamical equation without greatly affecting numerical performance, due to its simple dependence on the other dynamical equations.

## G Formulation of dynamics in terms of heritability *h*^2^(*t*) and mean fitness 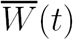

We work under the “closure” assumption of truncated Gram-Charlier moments and dynamical equations, as described above. The resulting two-variable dynamical system 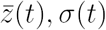 may instead be expressed in terms of 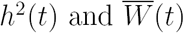, where

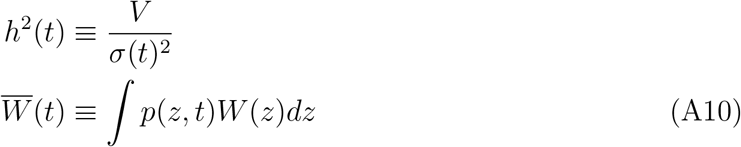

which have the dynamical equations

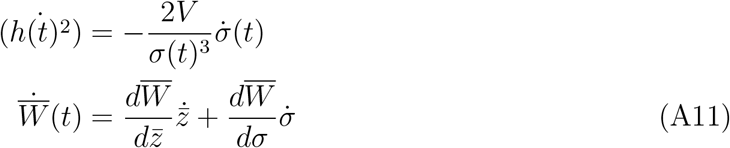

The exact form of the equivalence 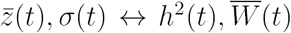 depends on the exact functional form of the fitness landscape *W*(*z*), but the general approach consists of the following steps:

### 1. Derive an expression for the fitness in terms of the trait moments

The fitness landscape *W*(*z*) is written as an infinite series in *z,*

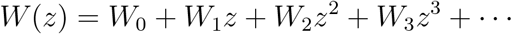

This series is substituted into Eq. A10, and then individual terms are averaged in order to generate a series representation of 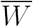 in terms of its raw moments 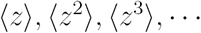 (as is also described in Appendix F). This series representation may then be converted into a sum over cumulants or moments using Eq. A3, and then the resulting series may be directly substituted into the dynamical equations for 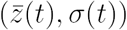. The resulting cumulant series may then be truncated in order to yield a closed form expression for 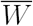. For clarity refer to this finite series expression as 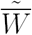.

### 2. Solve for the mean trait

The closed-form expression 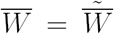 may then be algebraically inverted in order to generate a solution for the mean trait 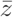 in terms of *σ* and 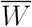. Importantly, because the mean appears in the explicit definitions of each of the raw moments given by Eq. A3, this step requires inverting a high-degree polynomial. Practically, symbolic solutions for 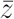 may be kept as placeholders until numerical values for individual parameters are chosen. We use the notation 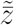 to refer to the values of the mean trait obtained by inversion of 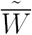

### 3. Substitute into the dynamical equations

We perform the following substitutions (in order) into the right hand side of the dynamical equations Eq. A11.

1. 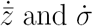 are replaced by their explicit expressions in terms of 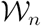 (as given in the main text)
2. Wherever it appears, 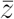 is replaced by 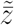, effectively replacing 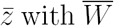 in the dynamical system.
3. Wherever it appears, *σ* is replaced by 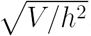, removing *σ* from the dynamical system.

For a specific functional form of *W*(*z*) (and thus 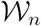), these steps may yield simple analytical results (as occurs for low-order polynomial or Gaussian fitness landscapes) or complex but complete analytic results, which require a symbolic algebra computer program to compute accurately.

## H 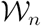 for exponential directional selection

A fitness landscape for exponential directional selection is given by

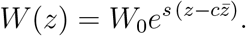

We insert this equation into the definition of 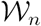,

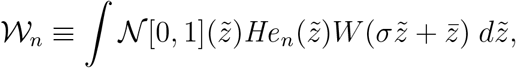

which results in a simple expression for 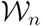 associated with exponential directional selection

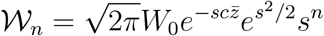

Inserting these terms into the truncated four-moment differential equations for 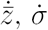,

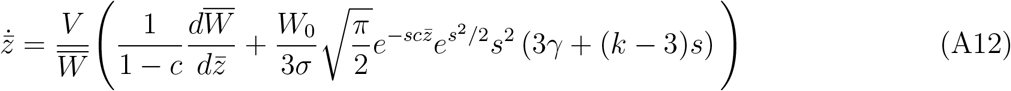

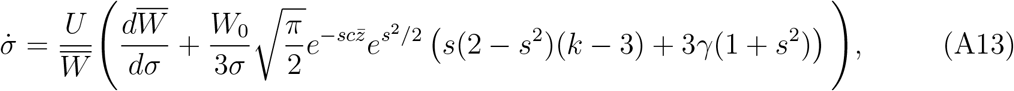

We note that similar equations are obtained by Bürger for a discrete-time model of cumulant dynamics under directional selection.^69^

Under the truncated model, we define the cryptic force of selection simply as the additional terms in each of Eq. A12 and Eq. A13, which would not be present if the trait distribution were purely Gaussian:

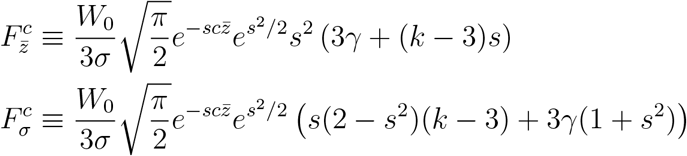

Depending on the overall strength of selection, these terms may be comparable in magnitude to the leading order gradient terms in the dynamical equations Eq. A12, Eq. A13.

## I Orthogonality of Hermite polynomials and shifted Hermite polynomials We wish to solve the following integral

We define a generating function,

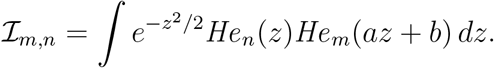

We define a generating function,

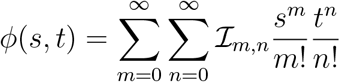

such that

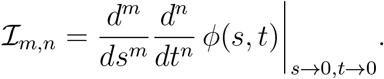

The generating function may thus be expressed as,

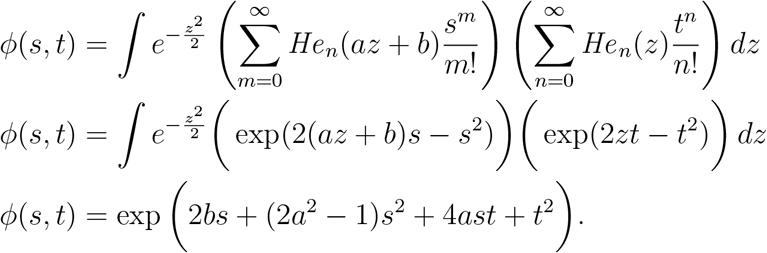

The derivatives of this generating function have a regular pattern due to the polynomial argument of the exponential, resulting in an analytic solution for the integral

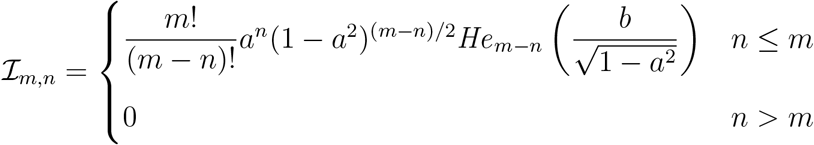

## J Orthogonality of Hermite polynomials and regular polynomials

Using the same method of characteristic functions used in Appendix I, it can be shown that the solution to the integral

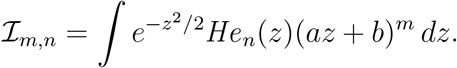

is

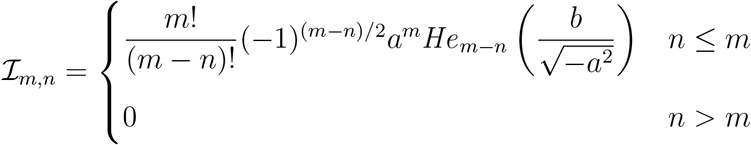

As an example application of this integral, we consider a fitness landscape given by a sum over Hermite polynomials

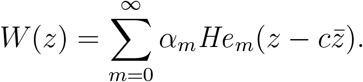

Inserting this expression into the definition of 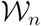 results in a series with fewer terms,

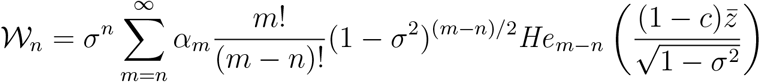

## K 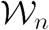 for polynomial fitness landscapes

We consider a fitness landscapes of the form

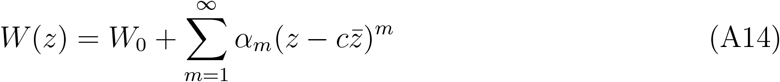

In general, analytic solutions are known for integrals of the form^38^

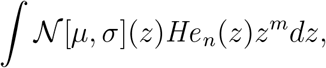

as given in Appendix J. Using these results, we find that the coupling coefficients 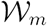 themselves have the form of an infinite series,

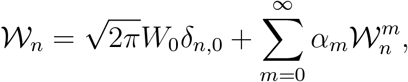

where

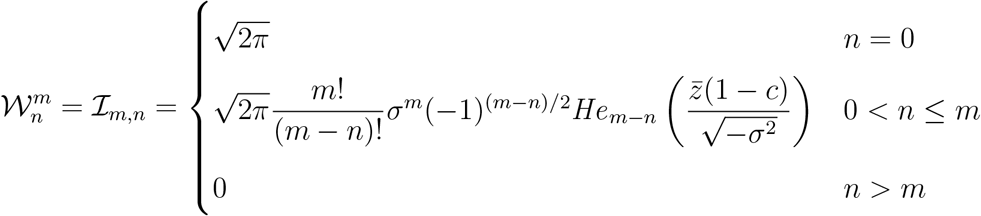

## L Comparison with other models of cumulant dynamics

Here, we describe the relationship between our approach and those taken in previous studies that have formulated natural selection in terms of moment or cumulant dynamics equations. We note that, to our knowledge, our specific model and derivation—inserting a Gram-Charlier series into the breeder’s equation, and then iteratively finding terms—is distinct from other approaches previously taken in existing literature. Our work is primarily relevant for describing phenotypic evolution over intervals in which the additive genetic variance *V* remains constant, such as during the phenotypic Wright-Fisher simulations we describe in the main text. The more detailed models described below include specific underlying genetic models; extending our results to work within these frameworks may require adding additional terms to the cumulant-wise generalized breeder’s equation Eq. A2, such as those found by Turelli and Barton 1994 for a broader study of non-Gaussian evolutionary processes.^23, 28^ However, the techniques that we use (the Gram Charlier series and cumulant-wise expansions of the fitness landscape) have also appeared in other contexts in evolutionary theory,^36^ such as the infinitesimal model in which a continuous genotype is associated with a continuous fitness landscape.^30, 67^ However, due to its origin in phenotypic theory, our approach entails certain distinct assumptions and regimes of applicability which we aim to outline below.

Zeng 1987 reports, to our knowledge, the earliest application of the Gram-Charlier series to a problem in population genetics.^48^ Zeng expands the joint distribution of genotypes and phenotypes in terms of a multivariate Gram-Charlier series (Eq. 3 of the referenced paper), and then studies the discrete-time consequences of truncation selection on an initially bivariate Gaussian distribution. Similar results are presented using different techniques in Bulmer 1980.^30^ Section 3.2.1 of Zeng provides the general form of cumulant iteration equations for the infinitesimal model under random mating, and Section 3.2.2 describes iteration equations under an explicit multilocus theory. For the multilocus case, his Eq. 46, 47 represent explicit dynamical equations for the mean and variance under truncation selection. The other results in the paper require explicit cross terms that depend on the fitness landscape to be calculated, a landscape-dependent calculation similar in principal to the 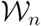 coefficients we describe above.

Related to the notion of dynamical equations arising from a cumulant-based approximation of the fitness distribution, Barton and Turelli 1987 presents a continuous-time matrix equation for the temporal dynamics of individual moments.^47^ The individual terms in the symmetric transfer matrix have a form in which the *i*^*th*^ term in a given row or column depends on the mean allelic effect of the *i* + 1^*th*^ and lower moments. They then show show for certain assumptions (such as diallelic inheritance) this matrix equation becomes finite, allowing the dynamics to be analyzed in closed form. The authors successfully use this model to describe the well-known mutation-selection balance effect.

Turelli and Barton 1990 extend this model in order to include recombination and linkage effects.^40^ The theoretical framework of their model focuses on the distribution of genotypes, each of which is assumed to be associated with a single phenotype—making this work comparable to the continuous trait approach we present in our work above. Within a multilocus framework that includes linkage, Turelli and Barton present discrete time iteration equations in terms of infinite summations over gradients of the mean fitness 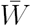 with respect to various phenotypic moments, each multiplied by its respective phenotypic moment. These recursions simplify under the assumption of many loci (the infinitesimal model), resulting in a linear “variance heritability” equation (Eq. 3.2b of the referenced paper) similar to the one found by Bulmer 1971.^28^ While our model above encodes information about the fitness landscape in terms of a set of “coupling coefficients” 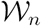, under the framework of Turelli and Barton 1990 information about the fitness landscape is encoded by computing the mean fitness gradients with respect to the moments—making these gradients the primary determinants of the tractability of a given landscape. We compare the mean trait evolution equation found by Turelli and Barton 1990 in the many-loci limit (Eq. 3.15 of the referenced paper) to the dynamical equations we derive for 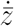 above. For a directional landscape, our results have slightly different coefficients (arising primarily from the continuous-time derivation and the form of our generalized breeder’s equation Eq. A2). However, the form of our solutions for 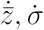 qualitatively match Eq. 3. 23 of Turelli and Barton (when higher order selection gradients are added back into their expression); in particular, the lowest order term is pro-portional to the additive genetic variance. The qualitative stability properties of Turelli and Barton’s results for a Gaussian fitness landscape entails global convergence for a stabilizing landscape, similar to our results for a polynomial stabilizing landscape.

Bürger 1991 notes the relative tractability of using cumulants, as opposed to moments, in the dynamical equations, and introduces the notion of taking a series expansion in the fitness landscape (Eq. 3.3 of the referenced work) in order to provide iterative terms in the dynamical equations (Eq. 4.8a,b).^37^ These equations have a similar form to Eq. 3.13 of Turelli and Barton 1990,^40^ although the latter is expressed in terms of moments. Bürger notes that linear selection advantages in a standard discrete time fitness model correspond to a Malthusian fitness landscape; we emphasize that due to the continuous time dynamics we study in the model we present above, the equivalent to this landscape in our formalism is simply a linear (and not exponential) function. Otherwise, the general dependence of terms in Bürger’s directional selection equation on the strength of directional selection has a power series form similar to the one we derive above.

Bürger 1993 studies cumulant dynamics under both directional and stabilizing (Gaussian) fitness landscapes.^69^ For the directional terms, the drift of an initially Gaussian distribution away from Gaussianity due to selection and mutation has the form of a power series in the selection strength multiplied by a Gaussian (Eq. 16 of the referenced paper); we note that these results are similar to those obtained for the directional linear and exponential landscapes above, with the linear directional landscape in continuous time being most analogous to linear directional selection in discrete time. Bürger also notes the timescale isolation associated with the dynamics of different cumulants. In the model we present above, this timescale isolation is used as an *ad hoc* heuristic to justify truncating the series of differential equations, since a finite number of differential equations may describe the dynamics for short timescales. Bürger also presents numerical results via Monte Carlo simulations illustrating the dynamics of cumulants in different fitness landscapes, and finds that the skewness and kurtosis generally remain bounded over a range of selection strengths. The coupling between selection strength, skewness, and the time evolution of the variance in these results is consistent with Barton and Turelli 1987,^47^ as well as the calculations we present above for a directional fitness landscape, which suggest a strong role of tail effects in determining the dynamical regime.

Barton and Turelli 1994 start from the breeder’s equation and develop a general non-Gaussian infinitesimal model.^23^ Their formalism takes root in the concept of using convolutional integrals to construct discrete-time integral iteration equations, a method originally developed by Slatkin 1970.^67^ Assuming a Gaussian transmission kernel centered on the midparent breeding value, Turelli and Barton note that cumulant iterations are a natural simplification for such integrals due to the linearity of cumulants under convolution—an approach that motivates our use of them as well above, and which is also discussed by Bulmer 1980.^30^ However, as with the method that we present above, Turelli and Barton note that cumulants present issues when recombination is included, due to a multiplicative increase in the number of cross-cumulant terms in the dynamical equations (Eq. 13 of the referenced work). As in Turelli and Barton 1990, they note that the primary challenge in constructing cumulant iteration equations for a given fitness landscape consists of computing closed form expressions for the gradients of the mean fitness with respect to individual cumulants. Analogously to the “coupling constants” 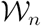 that we describe above, Turelli and Barton present an integral equation (Eq. 26 of the referenced paper) that allows individual terms of the fitness landscape to be included iteratively in the dynamics. They calculate these gradients analytically for the case of truncation selection, in which case the cumulant series readily truncates due to the leading order effects of truncation acting primarily on lower moments—resulting in dynamical equations complementary to those found by Zeng 1987.^48^ Turelli and Barton also provide general cumulant iteration equations in terms of selection gradients for both an explicit multilocus model (Eq. 35-40 of the referenced paper) and for the limit of an infinitesimal model, in which an arbitrarily large number of alleles may contribute to the fitness (Eq. 42, 43). Because the formulation of our phenotypic evolution model is mathematically similar to an infinitesimal genetic model, the dynamical equations Eq. 42, 43 of Turelli and Barton have a comparable form to the general dynamical equations for 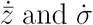 we calculate. However we truncate our series based explicitly on the order of the cumulant, whereas the selection gradient approach may allow higher cumulants to appear due to residual terms in a given derivative of the mean fitness 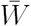. Turelli and Barton find that their iteration equations reduce to a higher-order variant of the breeder’s equation under the assumption of Gaussian breeding value distributions (Eq. 45). We note that we have assumed breeder’s equation type construction for the underlying genetics of our phenotypic model; an extension of our approach with a specific underlying genetic process may require additional terms to be added to the breeder’s equation.

Distinct from the approaches used above, Prügel-Bennet studies explicitly the natural selection process as an iterated random sampling process biased by the fitness, an approach similar in motivation to the phenotypic evolutionary model we derive above.^42, 50, 70^ Prügel-Bennet’s formulation of the selection process is parametrized by a selection rate parameter, *β*, analogous to the inverse temperature of a thermodynamic ensemble. Analytic forms of the cumulant dynamical equations require taking the equivalent of a “high temperature limit,” wherein the sampling exponential is linearized and thus extremal phenotypes are sampled less frequently. Our model presented above is similar to a continuous time, continuous trait formulation of this approach, with individual coupling coefficients 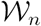 analogous to the individual series terms computed by Prügel-Bennet for the case where some phenotypes have a linear fitness advantage.

Another cumulant-based approach for a discrete allele sampling process is provided by Rattray and Shapiro, who study a case of multiplicative selection in which the number of beneficial alleles accrued by an individual confers that individual with a linear fitness advantage relative to the rest of the population.^39^ This multiplicative fitness landscape in discrete time is equivalent to the linear directional fitness landscape we study above for continuous time. Rattray and Shapiro exploit a useful property of multinomial sampling processes: in general, the *n*^*th*^ cumulant of the distribution may be written as a summation over terms of order 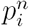 associated with the set of alleles indexed by *i* (Eq. 7 of the referenced paper). By assuming that each allele follows an independent diffusion process within the population, and that the growth rate of an allele in the diffusion equation is associated with its mean (the lowest cumulant), Rattray and Shapiro derive a series of dynamical equations describing the evolution of moments within the population (Eq. 12), which are linear (and thus may be solved via matrix inversion) for the multiplicative fitness landscape case. We note that our approach presented above differs in construction from Rattray and Shapiro, with the most substantive differences being (1) the explicit multilocus evolutionary model, which allows the cumulant equations to have forms as summations, and (2) the use of a diffusion process for individual alleles. In the many-allele limit implicit to the infinitesimal model, these conditions would describe circumstances with weak selection or extremely high mutation rates.

## M Numerical solution of cumulant dynamical equations

Efficient simulation of the dynamics of cumulants requires that moments and cumulants are not re-calculated at each time step, as integrals over the entire trait domain are generally computationally expensive to perform numerically. However, if high order cumulants are not heritable (and thus do not change in time), these values can be calculated in advance, thus truncating the number of quantities that must be calculate. Any distribution may be used to determine realistic constant values of these higher-order cumulants; in general, however, distributions are chosen that have cumulants that grow slowly in *n*—this is a requirement to safely truncate the Gram-Charlier series. Distributions used for the numerical work include the skew normal distribution, the exponentially-modified Gaussian distribution, and the uniform distribution.

The characteristic function for a given probability distribution is given by

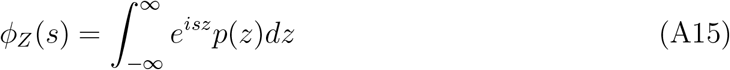

The cumulants are given by the derivatives of the logarithm of this distribution,

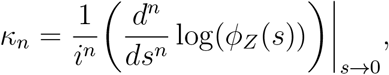

and they can generally be calculated to arbitrary order for a given distribution, abetting their use in numerical work. However, for certain distributions the series *κ*_*n*_ has a closed form. For example, for many of the examples simulated in this paper, the uniform distribution is used because it has the property *κ*_*n*_ = *B_n_/n,* where *B*_*n*_ is the *n*^*th*^ Bernoulli number. As a result, simulations of 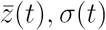 can be interpreted as the dynamics of a smoothed boxcar distribution with time-varying center and width. However, if other distributions with necessarily different cumulant series are used, the general dynamics will remain the same so long as the distributions and their associated cumulants result in a strictly decreasing series of magnitudes of Gram-Charlier coefficients |*c_n_|.* For numerical simulations shown in the paper, equivalent simulations were performed with a uniform distribution, a skew normal distribution, and an exponentially-modified Gaussian distribution, in order to ensure that qualitative results regarding the dynamics of the mean and variance remained the same.

For cases in which the underlying trait distribution as a function of time is needed, the time-dependent series of cumulant values, *κ*_*i*_(*t*), is first computed using the dynamical equations presented in the main text. These are then used to compute a time-dependent characteristic function,

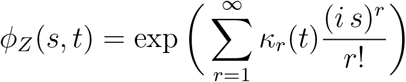

This expression derived by inserting a general Gram-Charlier series (Eq. A4) into Eq. A15. From *ϕ*_*z*_(*s*, *t*), the trait distribution may then be computed by numerically performing the inverse integral of Eq. Al5,

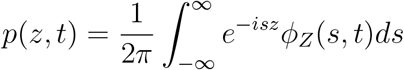

Using this equation, the dynamics of a sufficiently large number of cumulants may be converted to those of the full trait distribution. This underscores how moment transform methods are reminiscent of traditional spectral methods for dealing with partial differential equations, which provide insight into the properties of time evolution operators by recasting them within the frequency domain: similarly to Fourier modes, the moments of a distribution constitute parametrized integral transforms across real space. This analogy that can be made more direct when the distribution has a well-defined characteristic function, as it does here.

## N Numerical simulations of selection on individuals

Numerical simulations were performed using a variation of a standard Wright-Fisher process that accounts for selection on a continuous phenotype. For a given initial set of moments *µ*, *σ*^2^, γ, *κ*, a set of *N*_*i*_ phenotypes was initialized using Johnson’s *S*_*U*_-distribution (numerical optimization was used to select distribution parameters that produced desired moments). From these values, an initial mean trait value 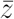 was calculated, allowing the full set of *z* values to be assigned relative fitnesses 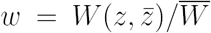. At the next timestep, *N*_*i*_ new values were drawn (with replacement) from the initial set of *z* values, and these values were taken as the “offspring” generation of phenotypes. This process was repeated for subsequent timesteps, with each offspring generation serving as the parent phenotype distribution for the next generation.

Because here we are primarily interested in the behavior of average moments of distributions in large populations, a relatively large value of *N_i_ =* 10^6^ was used in all simulations (effectively suppressing drift effects). However, smaller effective population sizes may be used in cases in which drift is important. In the simulations, the first four moments of the trait distribution vary gradually over time (primarily due to drift, here represented by the accumulation of sampling errors). However, as expected for large sample sizes, for simple fitness landscapes at large *N*_*i*_ the higher moments varied less than the lower moments over the simulation timescales, leading to a natural hierarchy in the timescales of dynamical variation in moments.

Figures S2-S4 show the results of Wright-Fisher simulations for the various fitness landscapes and conditions explored in the main text:

Figure S2 shows the dynamics of the mean trait 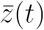 and trait range *σ*(*t*) under an exponential fitness landscape. The dynamics match those found for the simplified model in Figure 1; positive skewness produces gradual widening of the trait distribution, which in turn allows natural selection to proceed more rapidly when compared to the null case of a purely Gaussian distribution. Conversely, the case with negative skewness lags behind the default rate of natural selection, as predicted, resulting in a gradual narrowing of the trait range. For the timescales explored in these simulations, both the skewness *γ* and the kurtosis *k* are observed to fluctuate by less than 1%, despite there being no *a priori* constraints on their dynamics in the simulations. At smaller population sizes these quantities vary more due to drift, leading to more pronounced short-timescale departure from the the theoretical predictions.

Figure S3 shows the results of Wright-Fisher simulations in a quartic fitness landscape with all parameters equal to those in the disruptive selection panel shown in Figure 2. Due to constraints on the properties of Johnson’s *S*_*U*_-distribution, arbitrary values of *γ*, *k* cannot be probed over the full range of values shown in the Figure. Instead, we choose a line of values of *γ* and *k* that crosses through regions in which the intermediate phenotype equilibrium is active and inactive.We initialize each Wright-Fisher simulation at the each value of *γ*, *k* along this line, and set the initial mean and standard deviation to match the equilibrium solution of the cumulant model at that particular value of *γ* and *k.* We then simulate the Wright-Fisher dynamics for a short time interval, in order to yield an estimate for the velocity of the mean and trait range 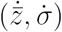 in the vicinity of the equilibrium point. These velocity values are difficult to directly compare to the eigenvalues yielded by the local stability analysis in Figure 2; instead, we convert these velocity values into an “angle of motion” towards or away from the equilibrium using the transformation *θ_sim_ =* arctan(*v*_*y*_/*v*_*x*_). We then compare this angle to the direction of the eigenvector associated with the largest eigenvalue of the local stability matrix for the cumulant dynamics model, in order to assess whether the local dynamics of the Wright-Fisher system match those predicted by the local stability of the simpler cumulant dynamical model. Figure S3 shows the resulting comparison; at large values of γ, the two cases nearly exactly coincide. This strong coincidence is expected due to the extremely short timescales over which local stability analysis predicts the dynamics, because drift in the values of the individual cumulants does not have time to substantially affect the relative dynamics.

Figure S4 shows the result of numerical simulations of for identical parameter values to those used in Figure 3. To facilitate easier comparison, a panel from that Figure is reprinted adjacent to the numerical results. It is apparent that the numerical work shows the same general trends as the theoretical model, including differences in the growth of the heritability depending on the sign of the skewness, and non-monotonicity in the dynamics that depends on the relative value of the kurtosis. For these simulations, the values of the skewness and kurtosis were observed to vary by less than 5% over the duration of the dynamics. Many of the qualitative differences in the dynamics (particularly at large values of the kurtosis *k*) occur due to the the accumulated effects of these moments gradually varying.

**Figure S1.**
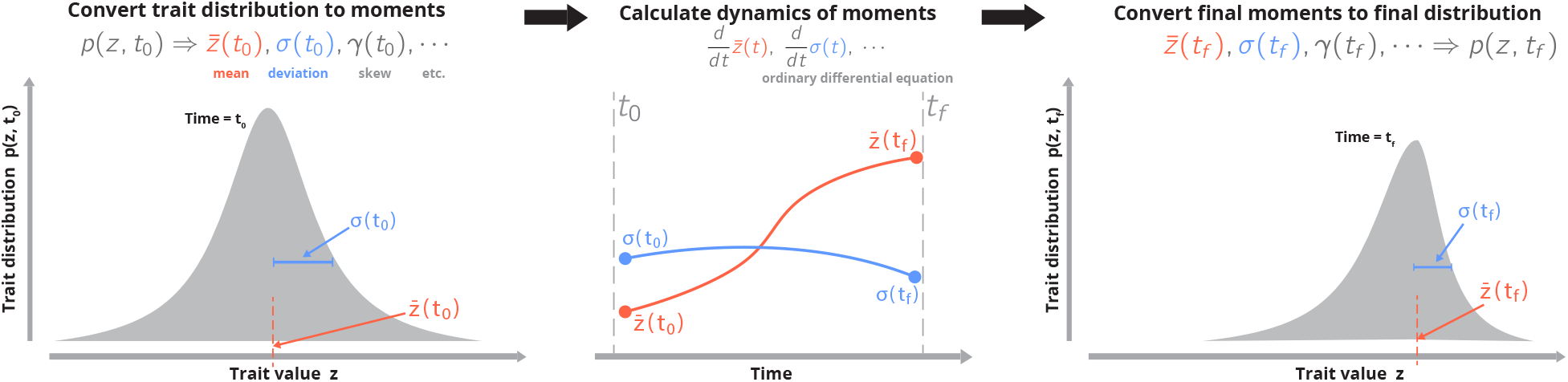
Overview of model approach. (Left) An initial distribution of traits in a population is parametrized into an arbitrary number of moments (the mean trait, trait range, skew, kurtosis, etc), which, collectively, describe the full state of the population at the initial time *t*_0_. (Middle) Using the breeder’s equation, a known fitness landscape, and a series expansion of the trait distribution, a set of ordinary differential equations is derived that describes the time-evolution of each moment of the trait distribution, with the moment values in the initial trait distribution acting as initial conditions. (Right) The solutions of these differential equations at some later time may then be used to reconstruct the full trait distribution at that time.

**Figure S2.**
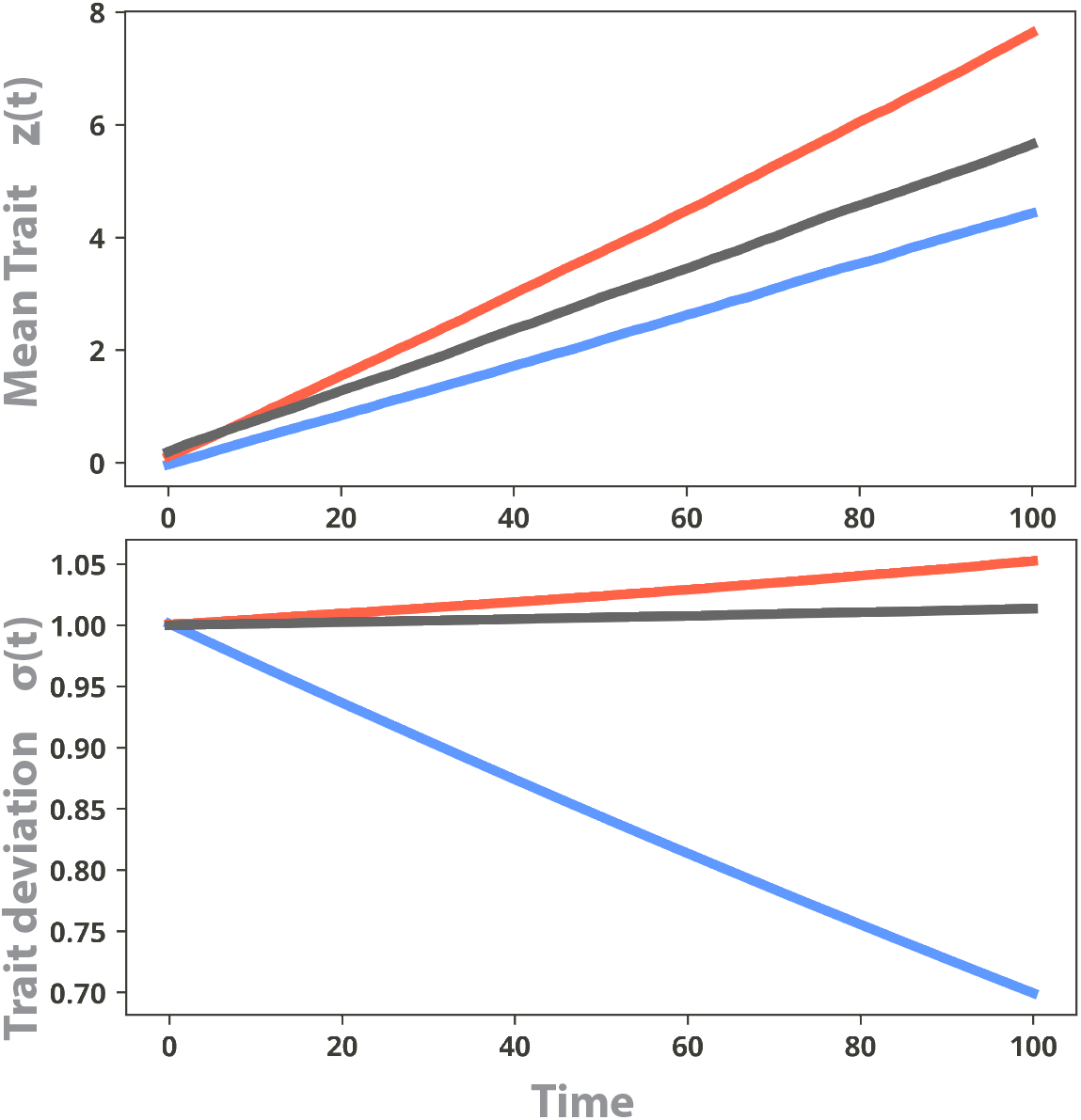
Wright-Fisher dynamics in exponential fitness landscape. The mean trait value and trait range for Wright-Fisher simulations in an exponential fitness landscape. Parameters are chosen to match those of Figure 1: gray curves correspond to an initially Gaussian population, blue curves correspond to initial distribution with negative skewness, and red curves correspond to initial distribution with positive skewness.

**Figure S3.**
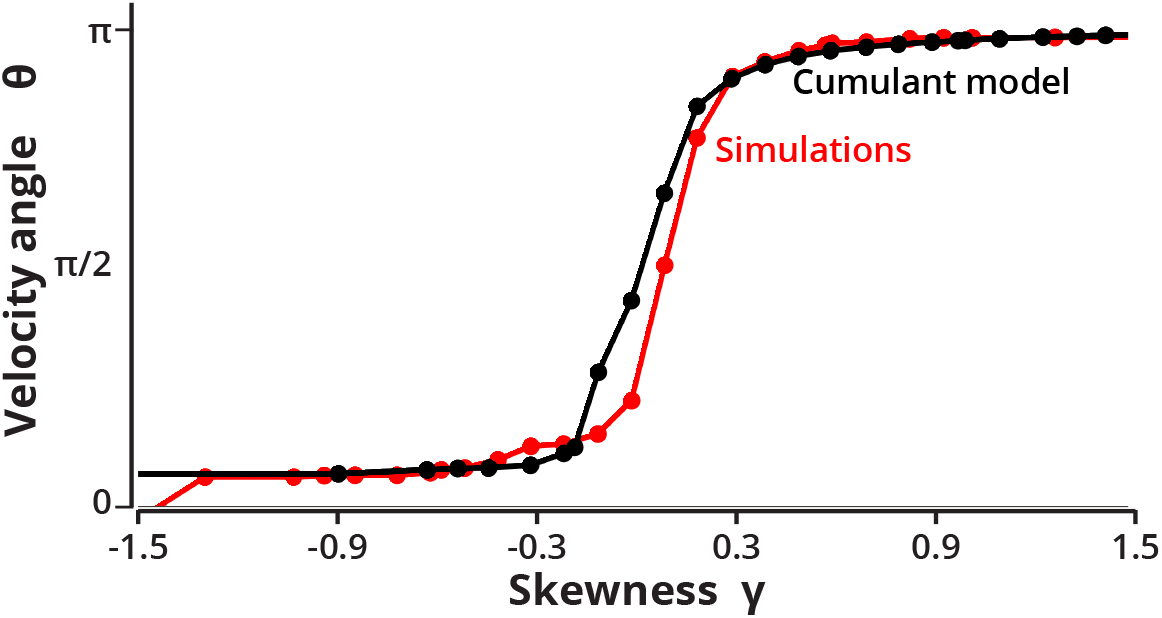
Wright-Fisher dynamics in quartic fitness landscape. A variety of *γ* values was first chosen, followed by a range of corresponding *k* values along the line *k = γ*^2^ + 2. At each point, the cumulant dynamical equations under the four-moment model were solved and the direction of the principal eigenvector associated with the dynamics was computed (black traces). For each value of *γ, k,* corresponding values of 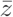 corresponding to the equilibrium point associated with the four-moment model were then used as starting distributions for Wright-Fisher dynamics simulations. From the short-time dynamics of the simulations, the direction of travel relative to the equilibrium point is calculated and shown in red on the plot.

**Figure S4.**
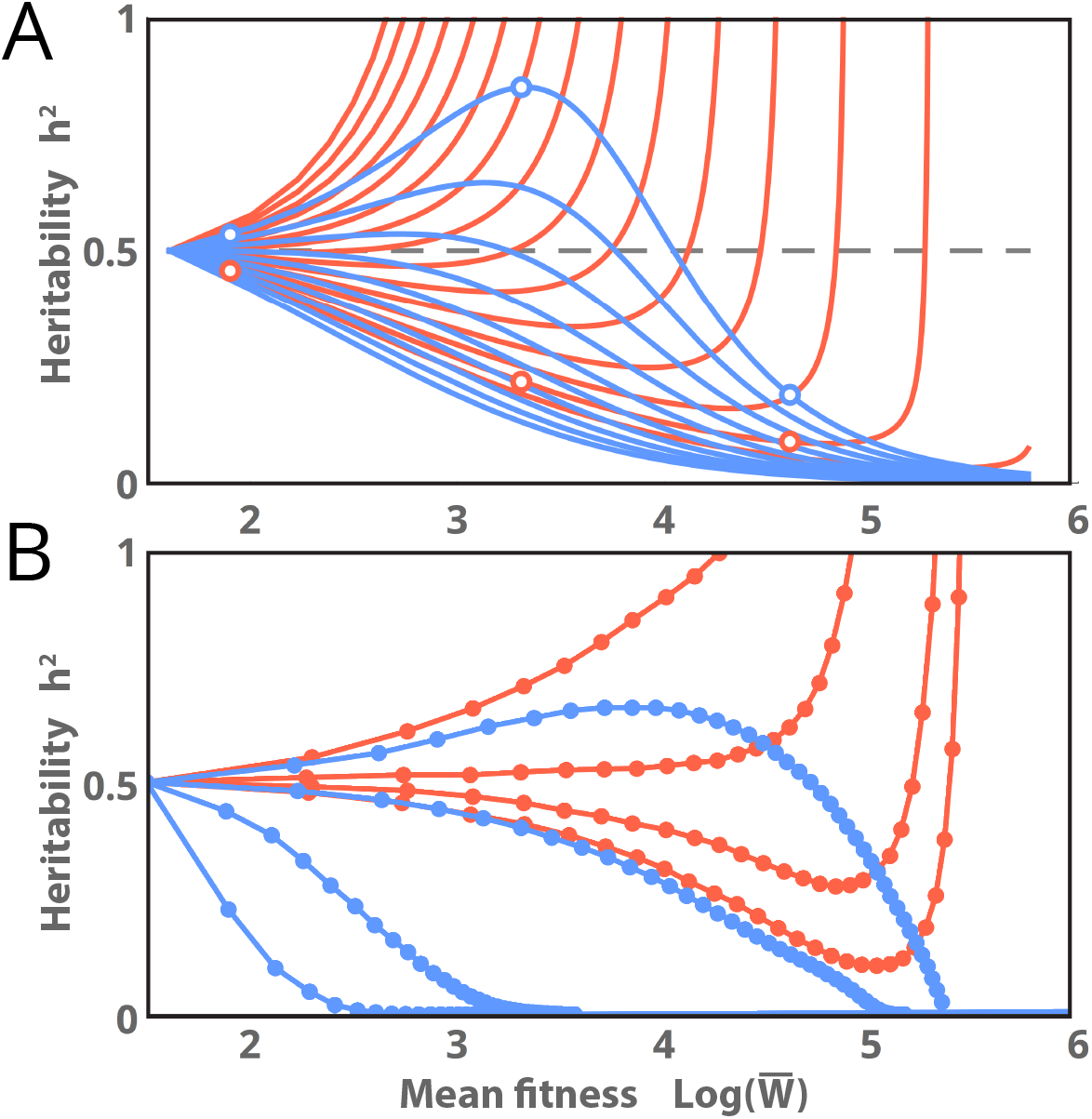
Comparison of analytic work with individual simulations. (A) Analytic results for heritability vs mean fitness under the four-moment model, with a linear fitness landscape. This panel is identical to Figure 3A of the main text. (B) Results of a numerical simulation of the Wright-Fisher model with 10^6^ individuals in the same linear fitness landscape of (A); blue traces correspond to initial conditions with *γ* = 0.05 (blue), red traces correspond to initial conditions with *γ* = *−*0.05, and lower traces within each colored group correspond to linearly increasing initial values of the kurtosis *k* between *γ*^2^ + 1 and *γ*^2^ + 6. All parameter values are constant, with values as given in Figure 3

